# Quantitative MRI Preprocessing: Effects of Tissue-Specific Smoothing Approaches on Statistical inference

**DOI:** 10.64898/2026.08.24.746651

**Authors:** Antoine Jacquemin, Christophe Phillips

## Abstract

**Background:** Quantitative MRI (qMRI) provides voxel-wise measurements of tissue properties related to myelin, iron and water content, making it a powerful tool for studying brain aging and microstructural alterations in vivo. However, conventional spatial smoothing can introduce partial-volume effects and blur tissue boundaries, potentially affecting both statistical sensitivity and anatomical specificity. Several tissue-specific smoothing strategies have been proposed to address these limitations, yet their relative impact on voxelwise statistical analyses remains insufficiently characterized. The present study aims (i) to systematically compare three tissue-specific smoothing strategies: a linear tissue-weighted compensated approach (TWS), a generalized version of nonlinear tissue-masked compensated smoothing approach (gTSPOON), and an intensity-weighted edge-preserving approach based on the Smallest Univalue Segment Assimilating Nucleus smoothing (SUSANs), and (ii) to investigate how smoothing approaches interact with statistical inference frameworks by comparing parametric and non-parametric voxel-wise analyse.

**Methods:** Analyses were performed on a publicly available lifespan qMRI dataset comprising 138 healthy participants (19–75 years) and quantitative maps of MTsat, PD, R1, and R2*. The generalized TSPOON (gTSPOON) method was implemented using tissue-specific masks derived from probabilistic tissue segmentation. All three smoothing approaches (TWS, gTSPOON and SUSANs) were parameterized to achieve comparable nominal spatial smoothing. Age-related effects were investigated separately in GM and WM using voxel-wise general linear models following a previously published framework. Statistical inference was assessed using multiple complementary approaches, including parametric Random Field Theory (RFT), under both stationarity and non-stationarity assumptions, as well as non-parametric permutation-based inference. In addition to conventional thresholded statistical parametric maps, voxel-wise log-likelihood (LL) maps were computed to quantify general linear model (GLM) goodness-of-fit independently of statistical thresholding. Bland–Altman analyses and spatial agreement metrics were subsequently used to compare smoothing strategies.

**Results:** TWS and gTSPOON produced highly similar spatial distributions of age-related effects across all qMRI parameters and tissue classes. However, TWS consistently yielded a larger number of significant voxels and clusters, reflecting slightly higher sensitivity, from slightly wider effective smoothness and reduced RESEL counts. By contrast, SUSANs generated substantially fewer significant voxels and clusters, associated with approximately half the effective smoothness and a markedly larger number of RESELs. Despite these differences in statistical sensitivity, voxel-wise LL analyses revealed distinct anatomical preferences for each smoothing strategy. TWS provided the best model fit predominantly within GM, whereas gTSPOON showed superior performance in homogeneous WM regions. Conversely, SUSANs achieved the highest LL values at GM–WM interfaces, particularly within sulcal and gyral transitions, indicating improved preservation of sharp anatomical gradients. These spatial patterns were consistently observed across MTsat, PD, R1 and R2* maps. Comparisons across stationary and non-stationary RFT assumptions revealed only minor differences, while non-parametric inference produced highly concordant results, indicating that the primary source of variability originated from the smoothing procedure itself rather than the inference framework.

**Conclusions:** Tissue-specific smoothing strategies substantially influence both statistical sensitivity and voxel-wise model fitting in qMRI analyses. While TWS and gTSPOON provide highly consistent results, the edge-preserving SUSANs approach preferentially enhances model fit at tissue boundaries. Importantly, voxel-wise log-likelihood mapping revealed that no smoothing strategy is uniformly optimal throughout the brain; instead, each method exhibits anatomically preferential regions where model fit is maximized. These findings suggest that smoothing should be viewed as a region-dependent optimization problem and highlight voxel-wise LL mapping as a principled framework for selecting or developing adaptive smoothing strategies tailored to specific neuroanatomical structures and biological processes, including age-related brain changes.

## 1 Introduction

### 1.1 Quantitative MRI

Quantitative magnetic resonance imaging (qMRI) shifts brain MRI from qualitative contrast toward standardized measurement of tissue properties in physical units. Unlike conventional weighted MRI, whose contrast depends on both tissue characteristics and acquisition settings, qMRI enables more reproducible comparison across subjects, sites, and time points, and improves sensitivity to microstructural features such as myelin and iron [27, 34]. This makes qMRI particularly attractive for longitudinal and multicenter studies, where conventional T1-weighted imaging remains vulnerable to inter-site variability [27].

At the core of qMRI are biophysical models that correct for instrumental biases and estimate (semi-)quantitative intrinsic parameters including longitudinal relaxation rate (R1), effective transverse relaxation rate (R2*), proton density (PD), and magnetization transfer saturation (MTsat). Multi-parameter mapping protocols provide these measures in a standardized framework and have shown good reproducibility across sites and scanners, supporting their use for morphometric, quantitative and disease-oriented studies [35, 30]. Open-source implementations such as the hMRI toolbox have further lowered the barrier to acquisition, processing, and harmonized analysis [32, 3].

Beyond methodological standardization, qMRI has clear translational and neuroscientific value. In high-grade glioma, multiparameter hMRI detected recurrence-associated tissue changes weeks to months before overt recurrence on conventional MRI [29]. More broadly, qMRI has been applied to multiple sclerosis, epilepsy, aging, and other neurological conditions, where it can reveal microstructural alterations that are not visible on standard imaging [22, 6, 30]. In particular, voxel-wise quantitative MRI analyses have demonstrated widespread age-related differences in myelin-, iron-, and water-sensitive parameters across gray and white matter during healthy aging [6]. Remaining limitations are practical rather than conceptual: acquisition time, data processing complexity, and the need for robust harmonization across platforms still constrain routine deployment [29, 30]. Even so, qMRI is increasingly established as a framework for non-invasive tissue characterization and a promising bridge between structural imaging and in vivo histology.

### 1.2 Partial volume effect

Partial volume effects (PVE) constitute a major limitation in quantitative MRI, as in PET imaging [19], because the spatial resolution of MRI is insufficient to ensure that each voxel contains a single homogeneous tissue compartment. Instead, within the brain, most voxels contain varying proportions of gray matter (GM), white matter (WM), and cerebrospinal fluid (CSF), particularly near tissue interfaces and within highly folded cortical regions. Consequently, the measured qMRI signal reflects a mixture of tissue-specific properties weighted by their relative volume fractions rather than a pure microstructural measurement [33, 2]. This mixing effect introduces systematic biases in quantitative parameters sensitive to myelin, iron, or water content, thereby reducing anatomical specificity and obscuring subtle biological variations.

The impact of PVE is especially pronounced in voxel-wise analyses, cortical morphometry, and longitudinal studies, where small changes in tissue composition may be interpreted as genuine biological effects. In aging studies, for example, cortical thinning and ventricular enlargement further increase tissue mixing, making robust partial volume handling essential for accurate interpretation of qMRI metrics. Partial volume contamination may also propagate through preprocessing steps such as image resampling and smoothing, amplifying signal leakage across tissue boundaries and altering the statistical properties of the data.

To mitigate these effects, several correction and estimation strategies have been proposed. Model-based approaches incorporating tissue probability fractions can explicitly account for mixed tissue composition and improve the accuracy of quantitative estimates [2]. More recent standardized qMRI processing frameworks additionally emphasize harmonized segmentation, careful spatial registration, and tissue-aware preprocessing pipelines to improve reproducibility and cross-site consistency [28]. Overall, controlling partial volume effects is not merely a technical refinement but a prerequisite for reliable tissue characterization and biologically meaningful interpretation in quantitative MRI studies.

### 1.3 Smoothing approaches

Spatial smoothing is a fundamental preprocessing step in neuroimaging analyses. Its primary objective is to reduce high-frequency noise and improve the signal-to-noise ratio (SNR) by averaging neighboring voxel intensities. Beyond this denoising effect, smoothing also facilitates spatial alignment across subjects, by reducing anatomical variability, and contributes to the validity and sensitivity of voxel-wise multi-subject statistical inference by increasing the spatial coherence of the data. However, because smoothing inherently combines information from adjacent voxels, it may also alter the anatomical specificity of quantitative measurements. In quantitative MRI, where voxel intensities are directly related to underlying tissue properties, the balance between noise reduction and preservation of biological information is particularly critical. Understanding the limitations of conventional smoothing approaches and the motivation for tissue-specific alternatives therefore requires first considering the influence of partial volume effects.

#### 1.3.1 Gaussian smoothing

Gaussian smoothing (GS) is a standard preprocessing technique widely used in neuroimaging to reduce high-frequency noise, improve signal-to-noise ratio (SNR), and increase the validity of voxel-wise statistical inference [38]. The method consists of convolving the image with a three-dimensional Gaussian kernel, producing a weighted local average in which neighbouring voxels contribute according to their spatial distance from the center voxel [1].

GS is implemented in most major neuroimaging software packages, including SPM [36], FSL [18], and AFNI [11]. In voxel-based neuroimaging, GS plays a central role in satisfying the assumptions of Gaussian random field theory (RFT) used for multiple-comparison correction in statistical parametric mapping. However, because the Gaussian kernel is purely spatial and intensity-independent, smoothing inevitably mixes signals across anatomical boundaries. In quantitative MRI, where voxel intensities carry direct biological meaning, any smoothing may therefore increase partial volume contamination and reduce sensitivity to fine microstructural variations, particularly near GM–WM interfaces and thin cortical structures.

#### 1.3.2 Tissue-specific smoothing approaches

Several tissue-specific smoothing strategies have been proposed to preserve tissue integrity while maintaining the statistical benefits of spatial smoothing.

Tissue-weighted smoothing (TWS) was introduced in the voxel-based quantification (VBQ) framework, later reframed in the hMRI toolbox [3], proposed by Draganski et al. [13], and this same VBQ/hMRI approach was used in multiparameter mapping studies [35, 6]. In this approach, quantitative maps are multiplied by tissue probability maps before smoothing and subsequently normalized by the smoothed tissue weights. This procedure restricts signal averaging predominantly within the same tissue class and reduces contamination across them. By incorporating tissue probabilities into the smoothing kernel, TWS preserves quantitative contrast while remaining compatible with RFT assumptions commonly used in voxel-wise statistical analysis [6].

The TSPOON (Tissue-SPecific, smOOthing-compeNsated) approach relies on binary or thresholded white matter mask to constrain smoothing more strictly within WM [21]. Compared with probabilistic tissue-weighted methods, as in TWS, binary masking strategies reduce cross-tissue signal leakage and preserve sharper anatomical boundaries, particularly within homogeneous white matter regions.

Finally, non-linear edge-preserving approaches such as the Smallest Univalue Segment Assimilating Nucleus (SUSAN) have been applied to quantitative MRI preprocessing [31, 9]. Unlike conventional Gaussian or tissue-weighted smoothing, SUSAN defines smoothing weights jointly from spatial proximity and local intensity similarity. This joint spatial–intensity formulation allows adaptive filtering that preserves sharp tissue transitions while reducing noise. Such approaches are particularly attractive for qMRI because they preserve fine structural details and minimize intensity blurring at gray matter/white matter interfaces. However, because SUSAN is intrinsically non-linear and highly sensitive to parameter selection, its impact on effective smoothness and statistical inference might differ substantially from conventional Gaussian-based approaches.

### 1.4 Objectives

The primary objective of this work is to evaluate how different smoothing strategies influence voxel-wise qMRI analyses. To this end, the effects of TWS, TSPOON (generalized to both GM and WM), and SUSAN smoothing approaches were systematically compared by a using previously published qMRI data [6].

The second objective is to investigate how smoothing strategies interact with statistical inference frameworks by comparing parametric and non-parametric voxel-wise analyses. Particular attention was given to how smoothing affects effective smoothness, statistical sensitivity, and cluster-wise inference.

## 2 Methods

The technical aspects of smoothing, classic and tissue-weighted, are first defined, then the data and statistical framework considered are described, and finally the comparison tools used to compare the different approaches are introduced.

### 2.1 Smoothing Approaches

Gaussian smoothing (GS) constitutes the reference framework upon which most neuroimaging smoothing approaches are built. GS consists of convolving quantitative map signal *s* warped into the standard space *ϕ* with a three-dimensional Gaussian kernel *g*, yielding the smoothed image at position *v*:

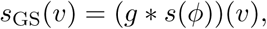

where denotes the convolution operator, and *v* is the spatial position vector in the standard space. The Gaussian kernel is defined as:

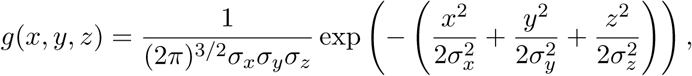

with *σ_x_*, *σ_y_*, and *σ_z_* denoting the Gaussian standard deviations along the three spatial dimensions, and (x,y,z)(x, y, z)(x,y,z) representing the coordinates of *v*. Through this convolution, each voxel intensity is replaced by a spatially weighted average of its neighbouring voxels, thereby reducing high-frequency noise and increasing spatial smoothness [14]. The extent of smoothing is commonly expressed by the “full width at half maximum” (FWHM) of the Gaussian kernel, which is related to the standard deviation by

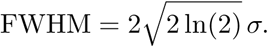

In the present work, three tissue-specific smoothing approaches will be investigated: TWS, gTSPOON and SUSANs.

#### 2.1.1 Tissue-Weighted Smoothing

The Tissue-Weighted Smoothing (TWS) method was first introduced by Draganski et al. [13]. For any voxel location *v*, TWS approach proposes to smooth quantitative map signal *s*, after spatial warping, for a specific tissue class (TC) density *ω* modulated for the deformation, as:

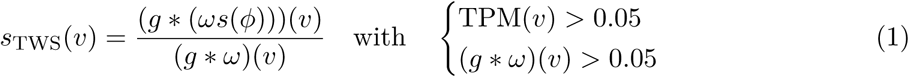

The numerator *g* (*w s*(*ϕ*)) in Equation 1 expresses the convolving of the quantitative MRI signal (*s*) weighted by (modulated) tissue class density (*ω*), both warped (*ϕ*) in a reference space, with an isotropic Gaussian kernel (*g*). The denominator simply consists in the smoothing (*g*) of the (modulated and warped) tissue class density (*ω*). Further masking is applied based on the thresholded (at *>* 0.05) prior tissue probability maps (TPM), and the smoothed (modulated and warped) tissue class density, i.e. the denominator in Equation 1. The rationale for the masking is two-fold: avoid signal spreading where there should a priori be none (TPM-based mask), and limit signal tissue-weighted smoothing where there is some actual tissue (*w*-based mask).

#### 2.1.2 Generalized Tissue-SPecific smOOthing compeNsated

Originally, the “Tissue-SPecific smOOthing compeNsated” (TSPOON) approach [21] was developed to take care of spatial smoothing in voxel-based analyses (VBA) of Diffusion Tensor Imaging (DTI) data, particularly in white matter (WM) regions. This approach is applied to DTI data, specifically focusing on fractional anisotropy and mean diffusivity measures derived from the eigenvalues of the diffusion tensor in WM.

Following the description of the original paper [21] and using the same notation as in Equation 1, the TSPOON operator can be described as:

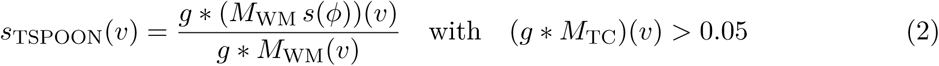

The numerator in Equation 2 expresses the convolving of the warped (*ϕ*) WM-masked (*M*_WM_) quantitative MRI signal (*s*) with an isotropic Gaussian kernel (*g*). The denominator simply consists in the smoothing (*g*∗) of the WM-masked *M*_WM_.

The generalization of TSPOON (gTSPOON) to both WM and GM requires the creation of both tissue-class–specific masks and, to avoid any overlap between these, we follow the same criteria as for qMRI-based publications [13, 6]. The individual smoothed, warped and modulated tissue probability maps, one for each intracranial tissue class (typically GM, WM and CSF), are averaged across all the subjects, providing a set of population-level smooth tissue probability maps. From these population-averaged maps, each voxel is then assigned to the tissue type with the highest probability, provided that this probability exceeds 20%, yielding binary masks for each tissue class (GM, WM, and CSF). The latter thresholding ensures specificity by excluding low-probability or ambiguous regions. Thus, gTSPOON can be expressed mathematically as follows:

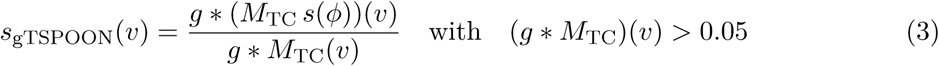

where *M*_TC_ in Equation 3 is the warped (*ϕ*) binary tissue-specific (either GM or WM) mask.

#### 2.1.3 Tissue-specific intensity-modulated smoothing with SUSAN

Originally introduced by Smith and Brady (1997) [31] and implemented in FSL toolbox [18], the “Smallest Univalue Segment Assimilating Nucleus” (SUSAN) approach preserves anatomical boundaries by restricting smoothing to voxels with similar intensities. For any voxel location *v*, the SUSAN smoothing (SUSANs) approach can be expressed as

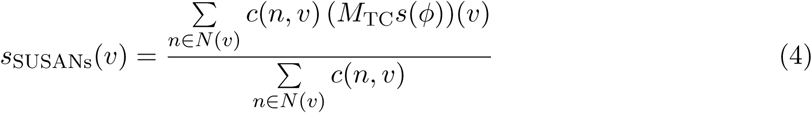

where *s*(*ϕ*) denotes the warped (*ϕ*) quantitative map signal *s* at voxel *v*. *c*(*n, v*) is the SUSAN combined weight integrating spatial proximity and signal intensity of each of the *n* voxels in the neighborhood *N* of the voxel at position *v*. This combined weight *c*(*n, x*) is expressed, for a intensity map *I*(*v*), as:

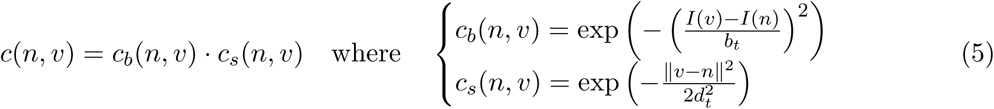

where *I*(*v*) and *I*(*n*) denote the quantitative MRI signal at voxel locations *v* and *n*, respectively, *b_t_* is the brightness threshold (controlling sensitivity to intensity differences and thus ensuring edge preservation), *d_t_* is the Gaussian spatial scale parameter (i.e. standard deviation in mm, 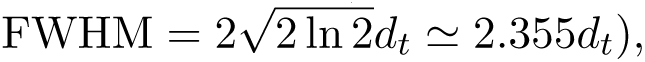 is the squared Euclidean distance in mm between voxels *v* and *n*.

The SUSAN weighting function combines spatial proximity and signal similarity. The spatial parameter *d_t_* determines the extent of the neighborhood over which smoothing is performed, whereas the brightness threshold *b_t_* controls the influence of neighboring voxels according to their signal similarity, thereby preserving anatomical boundaries while reducing noise.

### 2.2 Data and statistical framework

Quantitative data will be smoothed as described in the previous section, fed into different statistical models, then the obtained results will be compared. The FWHM kernel is isotropic with a width of 3 mm. To ensure a fair comparison with the Gaussian-based tissue-specific smoothing approaches, the SUSANs spatial parameter *d_t_* was chosen to produce an effective smoothing equivalent to a Gaussian kernel with a FWHM of 3 mm (*d_t_* ≈ 1.274 mm) and the brightness threshold *b_t_* was set to three-quarters of the median signal intensity of each quantitative MRI map, providing a data-driven balance between edge preservation and noise reduction.

#### 2.2.1 Dataset

The qMRI dataset used to illustrate this study was originally acquired and described by Callaghan et al. [6] in the context of investigating age-related microstructural changes in the healthy human brain. The dataset comprises MRI data from 138 healthy participants (89F/49M) ranging from 19 to 75 years of age (mean age: 46.6 years). This study assessed the impact of smoothing strategies on qMRI-based statistical analyses using modulated GM, WM, and CSF tissue-probability maps and normalized MTsat, PD, R1, and R2* quantitative maps in MNI space.

MRI acquisitions were performed on two 3T Siemens systems (*Trio* and *Quatro*) using a multiparameter mapping (MPM) protocol [35]. The spatially processed (i.e. quantitative maps reconstruction, and warping into standardized MNI space) dataset is publicly available, following the “Brain Imaging Data Structure” (BIDS) specifications [16, 20], through the OpenNeuro platform [7].

All qMRI maps were smoothed using TWS, gTSPOON and SUSANs. A target smoothing kernel of FWHM = 3 mm, as in [6] was converted into *σ* using the standard Gaussian relation *d_t_* = *σ* = ^FWHM^*/*8 ln(2). In SUSANs, the spatial smoothing parameter (*d_t_*) corresponds to the Gaussian standard deviation (*σ*) expressed in millimeters. For a FWHM of 3 mm, this yields *d_t_* ≈ 1.274 mm.

For SUSANs, modality-specific brightness thresholds (*b_t_*) were estimated from the quantitative MRI maps, from which voxels with inconsistent values (zero, negative, and NaN) were rejected. The SUSAN brightness threshold was then defined as: *b_t_* = 0.75 median(*I*_valid_) independently for each subject and modality.

#### 2.2.2 GLMs and inference

Voxel-wise statistical analyses were performed within the General Linear Model (GLM) framework implemented in SPM25 [36]. At each voxel, the observed quantitative values across subjects were modeled as *Y_v_* = *Xβ_v_* + *ε_v_*, where *Y_v_* denotes the voxel-wise observations, *X* the design matrix (containing the regressors of interest and nuisance covariates), *β_v_* the regression coefficients, and *ε_v_* the residual error term. Depending on the scientific question under investigation, both linear and non-linear model parameterizations can be specified through appropriate regressors within the design matrix.

To assess the robustness of statistical findings with respect to the inference framework, both parametric and non-parametric approaches were considered. Parametric inference was performed using Random Field Theory (RFT) as implemented in SPM [15, 38]. Classical RFT assumes that residual statistical fields are approximately Gaussian, sufficiently smooth, and spatially stationary. Under these assumptions, family-wise error (FWE) correction can be achieved through analytical approximations based on the geometry of the search volume and the estimated number of resolution elements (RESELs). Because preprocessing procedures may alter the spatial smoothness of the data, both stationary and non-stationary RFT corrections were considered. The latter accounts for spatially varying smoothness by incorporating local RESEL density estimates into the multiple-comparison correction procedure while preserving the underlying voxel-wise test statistics.

Non-parametric inference was performed using the “Statistical non-Parametric Mapping” (SnPM) toolbox [24, 37]. Unlike RFT-based approaches, permutation testing estimates the null distribution empirically through repeated permutations under the assumption of exchangeability. This framework avoids strong assumptions regarding residual normality, homoscedasticity, or spatial stationarity, and is therefore particularly suitable when preprocessing introduces heterogeneous smoothness patterns or non-linear transformations. Family-wise error control was obtained from the empirical distribution of the maximal statistic across permutations.

Together, these complementary inference frameworks enabled the evaluation of statistical results under different assumptions regarding residual distributions and spatial smoothness, providing a comprehensive assessment of the robustness of voxel-wise quantitative MRI analyses.

#### 2.2.3 Biological application

Although the statistical framework described above is applicable to a wide range of neuroimaging questions, a specific biological application is required to evaluate the influence of smoothing and inference strategies on quantitative MRI analyses. Brain aging constitutes a particularly relevant use case because it is one of the major sources of biological variability throughout adulthood and is associated with widespread microstructural alterations affecting myelin, iron, water content, and tissue organization [30]. Consequently, age-related qMRI analyses provide a well-established framework for assessing the sensitivity and robustness of voxel-wise statistical methods.

Importantly, age-related effects are not necessarily restricted to linear trajectories. While some quantitative MRI parameters exhibit approximately monotonic changes across adulthood, others may follow more complex patterns characterized by periods of maturation, stability, or accelerated decline [6, 23, 8]. Accounting for both linear and non-linear effects is therefore important for capturing the diversity of biological processes reflected by qMRI measurements.

For these reasons, the present study adopted the voxel-wise multiple-regression framework originally proposed by Callaghan et al. [6]. For each quantitative MRI parameter and tissue class, the primary model included age as the regressor of interest, while sex, total intracranial volume (TIV), and scanner type were incorporated as nuisance covariates. To investigate potential non-linear trajectories, an additional model including a quadratic age term (age^2^) was also evaluated. Continuous covariates were mean-centered prior to model estimation to reduce collinearity between linear and quadratic terms and improve parameter interpretability. Statistical inference was subsequently performed on contrasts testing with F-statistics, positive and negative associations with the age-related linear or non linear regressor.

Eventually, each of the 4 quantitative maps were thus smoothed in 3 different ways, separately for GM and WM. These 4 3 2 sets of smooth images were entered in 2 different GLMs, modeling linear age effect with or without an extra non-linear regressor, then 3 statistical inference procedures were applied (relying on stationary and non-stationary RFT, plus SnPM), to overall generate 144 statistical maps.

### 2.3 Comparison tools

To compare the effect of the different smoothing approaches on the resulting statistical maps, per quantitative map and tissue class, several tools will be employed to extend the scope of comparison.

#### 2.3.1 Summary statistics

In order to summarize statistical maps in a few numbers, we looked at the statistical threshold used to reach *p <.*05 FWER significance, and the number of voxels per RESEL, which is a proxy of the effective smoothing of the data. From the thresholded maps, we also collected the total counts of significant voxels and clusters,

#### 2.3.2 Bland-Altman Plot

Agreement between statistical maps was first assessed using Bland–Altman analysis [4, 5], a standard method for evaluating the consistency between two quantitative measurement techniques. Unlike correlation analysis, which only measures the strength of a linear association, Bland–Altman analysis evaluates the absolute agreement between paired observations and is therefore particularly suitable for comparing quantitative neuroimaging preprocessing strategies.

For two maps *S*_1_ and *S*_2_, the Bland–Altman representation is defined voxel-wise as

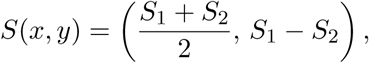

where the horizontal axis represents the mean of the paired measurements and the vertical axis their difference. This representation enables visualization of systematic biases, proportional deviations, and variability across the range of observed values.

In practice, the two volumetric images were vectorized after exclusion of invalid voxels, including NaN, infinite, and zero-valued entries. For each voxel pair, the mean intensity and the difference between methods were computed. The average difference corresponds to the bias between methods, while the standard deviation of the differences defines the limits of agreement, typically expressed as bias ± 1.96 × SD.

Within the context of qMRI smoothing comparisons, Bland–Altman analysis provides complementary information to statistical threshold maps by quantifying how preprocessing strategies modify the absolute quantitative values of tissue parameters. This is particularly relevant for tissue-specific smoothing methods, where preserving biologically meaningful qMRI intensities is a central objective.

#### 2.3.3 Voxel-wise log-likelihood

Voxel-wise log-likelihood (LL) maps were also computed to evaluate the quality of the GLM fit associated with each smoothing strategy [15, 25, 26]. Unlike binary statistical comparisons, LL maps provide a continuous model-based measure of goodness-of-fit and allow direct comparison between linear and non-linear smoothing approaches.

At each voxel *v*, the group-level GLM was defined as **Y**(*v*) = *X**β***(*v*) + ***ε***(*v*). Residuals were assumed to follow a Gaussian distribution ***ε***(*v*) (**0***, σ*^2^(*v*)**I**) with voxel-wise variance *σ*^2^(*v*) estimated from the residual mean squares image produced by SPM. Then for each subject *s*, the fitted signal was reconstructed as *µ_s_*(*v*) = *X_s_**β***(*v*) where *X_s,_*: denotes the *s*-th row of the design matrix. Under the Gaussian assumption, the voxel-wise log-likelihood is given by 6:

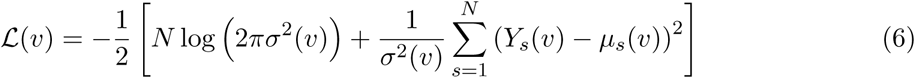

Higher LL (L) values indicate a better fit between the observed data and the statistical model, reflecting both lower residual error and lower estimated noise variance. In practice, LL maps were computed independently for each smoothing condition using the subject images, beta coefficients, and residual variance maps from the corresponding SPM models. Calculations were restricted to tissue-specific masks consistent with the statistical analyses. These maps enabled voxel-wise comparison of model fit across smoothing approaches and provided additional insight into how different smoothing strategies affect the statistical representation of tissue-specific qMRI aging effects.

#### 2.3.4 Similarity Metrics

Spatial agreement between thresholded statistical maps, obtained with different smoothing approaches, was quantified using the Jaccard Index (JI), Dice Coefficient (DC), and Cohen’s Kappa coefficient (*κ*). These complementary metrics evaluate overlap and agreement between binary voxel-wise maps [17, 12, 10]. Their mathematical definitions and interpretations are summarized in Table 1. Together, these metrics provide complementary information regarding the reproducibility, spatial consistency, and localization of age-related qMRI effects obtained with different smoothing strategies.

**Table 1:** Similarity metrics used to quantify spatial agreement between thresholded statistical maps.

| Metric | Equation | Interpretation |
| --- | --- | --- |
| Jaccard Index | $JI(A, B) = \frac{\vert A \cap B\vert }{\vert A \cup B\vert }$ | Measures the ratio between the intersection and union of two binary sets. Values range from 0 (no overlap) to 1 (perfect overlap) [17]. |
| Dice Coefficient | $DC(A, B) = \frac{2\vert A \cap B\vert }{\vert A\vert + \vert B\vert }$ | Quantifies spatial overlap while weighting the intersection more strongly than the Jaccard Index. Values range from (no overlap) to 1 (perfect overlap) [12]. |
| Cohen’s Kappa | $\kappa = \frac{p_o - p_e}{1 - p_e}$<br>with<br>$p_e = p_a p_b + (1 - p_a)(1 - p_b)$ | Measures agreement corrected for chance overlap. Values range from $-1$ (systematic disagreement) to 1 (perfect agreement), with 0 indicating chance-level agreement [10]. |

## 3 Results

### 3.1 Effect of smoothing on thresholded SPMs

Figure 1 illustrates representative voxel-wise threshloded statistical maps obtained for MTsat age-related effects, at a voxel-wise *p <.*05 FWER-corrected threshold using a SnPM inference, for the 3 smoothing approaches. Then Figure 2 shows the same results but only for SUSANs smoothing, with the prametric inference, with and without stationarity assumption.

**Figure 1:**
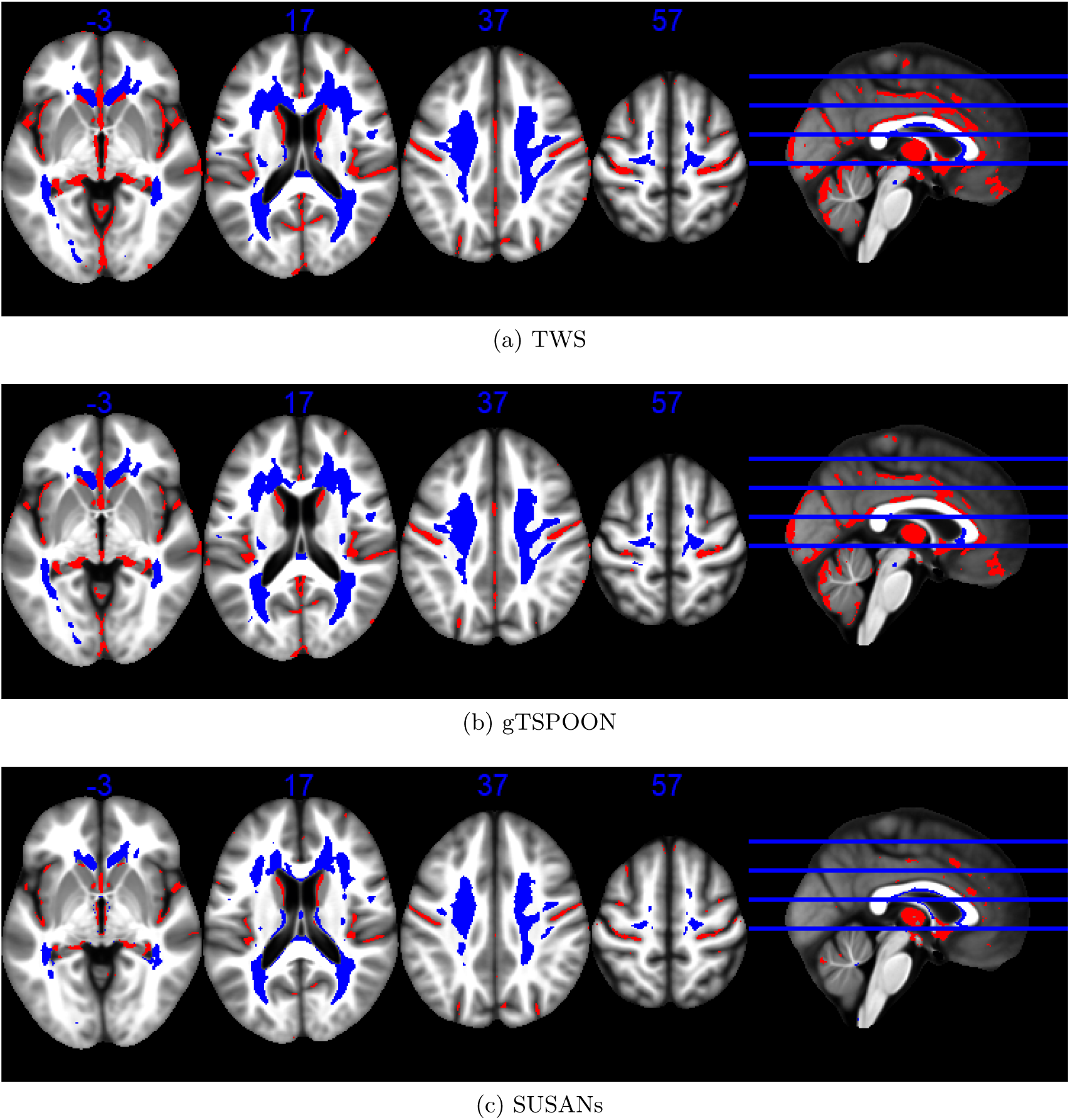
Statistical non parametric maps identifying regions (red for GM and blue for WM) in which MTsat significantly decreased with age at the *p <* 0.05 FWE corrected level using non parametric statistical inference (SnPM). The results are superimposed on the mean MT map for the cohort in MNI space. The four axial slices are located at z =-3, 17, 37 and 57 mm, from left to right, as illustrated on the sagital slice (right).

**Figure 2:**
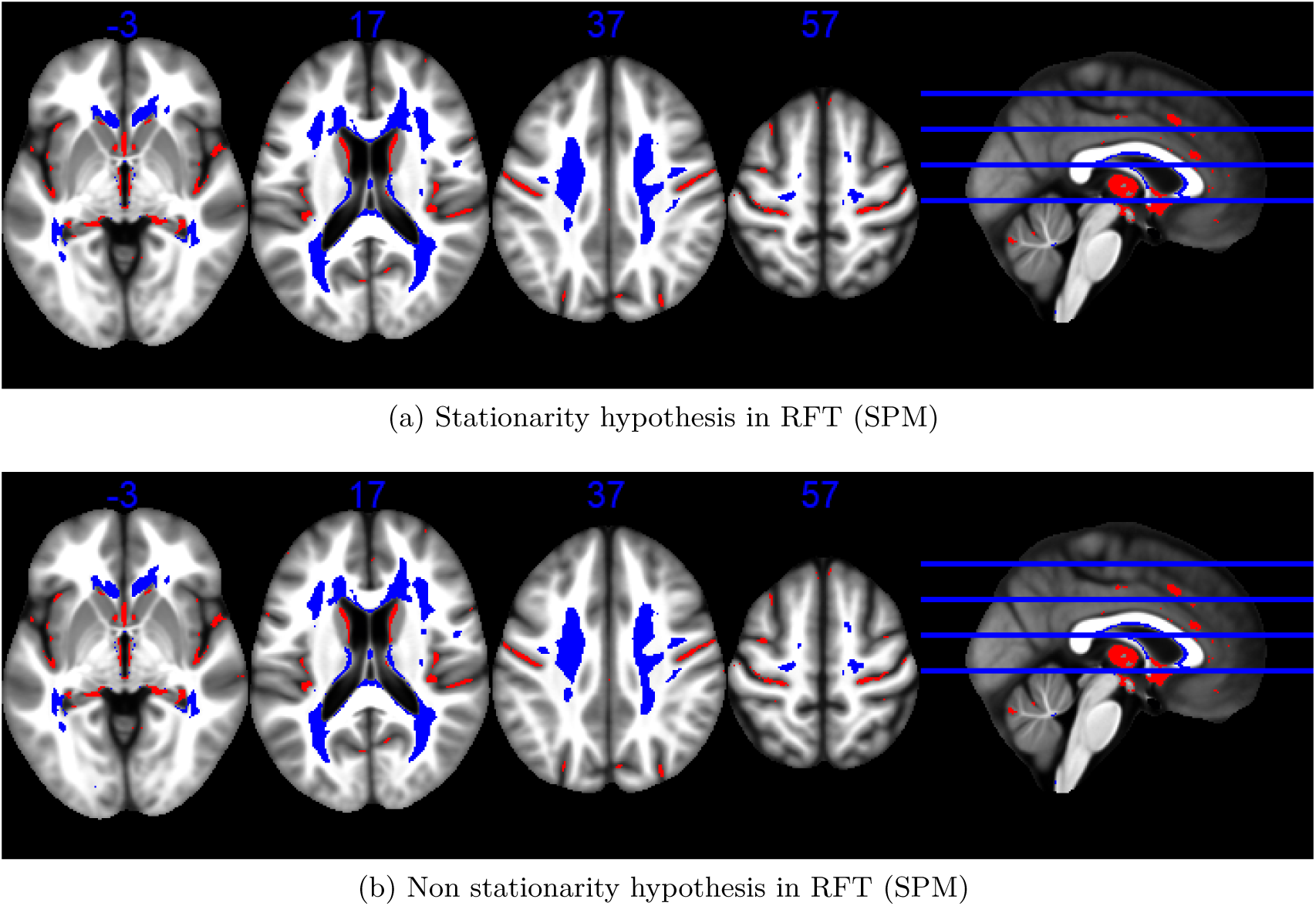
Statistical parametric maps identifying regions (red for GM and blue for WM) in which MTsat significantly decreased with age at the *p <* 0.05 FWE corrected level. The results are superimposed on the mean MT map for the cohort in MNI space. The four axial slices are located at z =-3, 17, 37 and 57 mm, from left to right, as illustrated on the sagital slice (right). SUSANs has been used for smoothing.

While all approaches revealed broadly similar spatial patterns of significant GM and WM voxels with MTsat values decreasing with age, there are some differences in statistical sensitivity, corrected thresholds, and effective spatial smoothness across smoothing, inference strategies, and qMRI maps. These parameters are further detailed in Tables 2 and 3.

**Table 2:** Overview of statistical T-thresholds values (*p <* 0.05 FWER) for different smoothing and statistical approaches across inference’s approaches, quantitative parameters and tissue types.

| Inference type | Smoothing | MTsat |  | PD |  | R1 |  | R2* |  |
| --- | --- | --- | --- | --- | --- | --- | --- | --- | --- |
|  |  | GM | WM | GM | WM | GM | WM | GM | WM |
| parametric under stationarity hypothesis (SPM) | TWS | 5,36 | 4,99 | 5,26 | 5,12 | 5,30 | 4,91 | 5,30 | 4,96 |
|  | gTSPOON | 5,38 | 5,01 | 5,27 | 5,13 | 5,32 | 4,93 | 5,30 | 4,95 |
|  | SUSANs | 5,64 | 5,38 | 5,64 | 5,53 | 5,64 | 5,39 | 5,64 | 5,46 |
| parametric under non stationarity hypothesis (SPM) | TWS | 5,36 | 4,99 | 5,26 | 5,13 | 5,30 | 4,91 | 5,30 | 4,96 |
|  | gTSPOON | 5,38 | 5,01 | 5,28 | 5,13 | 5,32 | 4,93 | 5,30 | 4,95 |
|  | SUSANs | 5,64 | 5,36 | 5,64 | 5,53 | 5,64 | 5,37 | 5,64 | 5,45 |
| non parametric (SnPM) | TWS | 5,18 | 4,67 | 5,04 | 4,99 | 5,14 | 4,72 | 5,06 | 4,79 |
|  | gTSPOON | 5,22 | 4,72 | 5,06 | 4,97 | 5,18 | 4,65 | 5,10 | 4,78 |
|  | SUSANs | 5,48 | 5,16 | 5,40 | 5,25 | 5,40 | 5,19 | 5,42 | 5,23 |

**Table 3:**
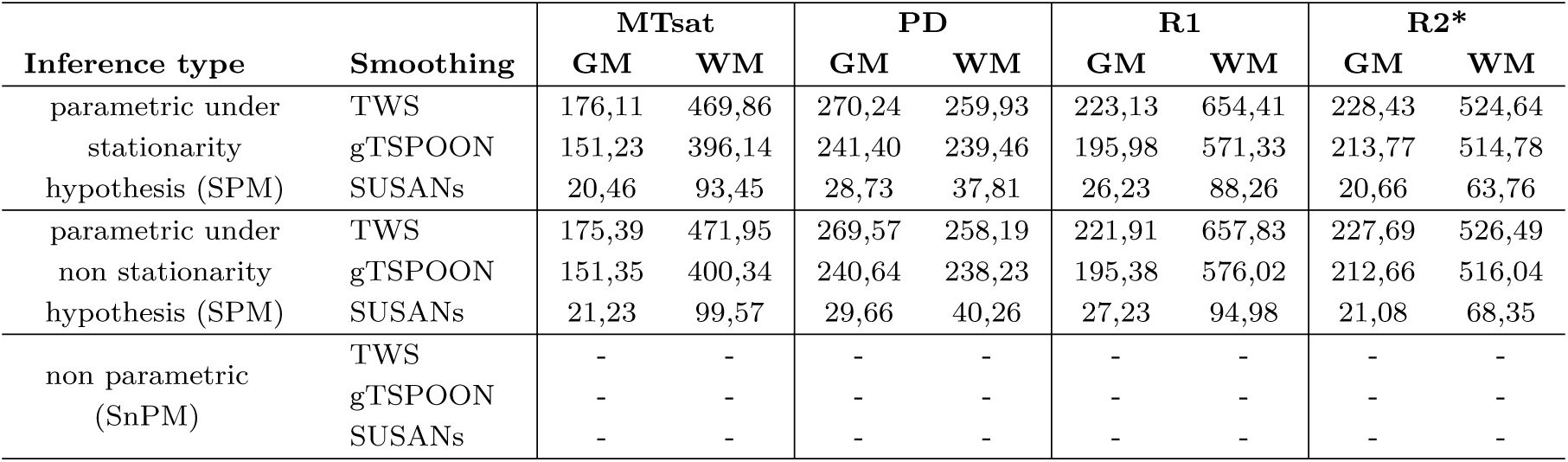
Overview of number of voxels per RESEL after inference (*p <* 0.05 FWER) for different smoothing and statistical approaches across inference’s approaches, quantitative parameters, and tissue types.

Table 2 summarizes the statistical T-thresholds corresponding to voxel-wise significance at *p <*0.05 FWER across smoothing strategies, inference frameworks, quantitative MRI parameters, and tissue classes. Across all inference approaches and qMRI maps, SUSANs systematically produced the highest statistical thresholds in both GM and WM, indicating more conservative voxel-wise significance criteria compared with TWS and gTSPOON. In contrast, TWS and gTSPOON yielded highly comparable thresholds, with only minimal differences across tissues and quantitative parameters. Under parametric inference assuming stationarity, T-thresholds for TWS and gTSPOON remained remarkably stable across qMRI maps, typically ranging between approximately 4.9 and 5.4. WM generally exhibited slightly lower thresholds than GM for these two smoothing approaches, particularly for MTsat, R1, and R2*. By comparison, SUSANs consistently showed elevated thresholds, reaching values up to 5.64 in GM across all qMRI maps, suggesting stronger smoothness-related corrections and reduced statistical sensitivity.

Similar results were obtained under parametric inference accounting for non-stationarity, with only negligible differences relative to the stationary framework. Although non-stationary random field theory models spatially varying smoothness, the representative corrected threshold reported by SPM was very similar to that obtained under the stationarity assumption. This observation suggests that incorporating non-stationarity had only a limited effect on the overall inferential calibration in the present dataset. However, this single reported value does not reflect the local spatial variability of the correction, which may differ across brain regions. As in the stationary analysis, TWS and gTSPOON yielded nearly identical representative threshold values, whereas SUSANs consistently produced higher values across tissues and qMRI maps.

TO DO: find a way to get the range of thresholds in the case of non stationarity assumption in RFT. Nothing in spm manual, through elicit or by looking at spm_uc. Looks that there is no good solution, either change some of spm scripts (without being sure of what we do, either our own script but strong hypothesis.

Under non-parametric inference (SnPM), all smoothing approaches exhibited reduced T-thresholds compared with parametric inference. TWS generally yielded the lowest thresholds overall, followed closely by gTSPOON, while SUSANs continued to produce systematically higher values. This reduction in threshold magnitude under SnPM suggests an increased sensitivity of permutation-based inference relative to parametric RFT approaches.

Table 3 reports the estimated number of voxels per RESEL after inference across smoothing strategies, inference frameworks, quantitative parameters, and tissue classes. Across all qMRI maps and tissues, TWS consistently produced the largest voxels-per-RESEL values, followed by gTSPOON, whereas SUSANs yielded substantially smaller values. These differences were especially pronounced in WM, where TWS reached values above 650 voxels per RESEL for R1 maps, compared with fewer than 100 voxels per RESEL for SUSANs.

The largest voxels-per-RESEL estimates were systematically observed in WM for TWS and gTSPOON, particularly for R1 and R2* maps, indicating a substantially smoother effective spatial structure. In contrast, SUSANs consistently exhibited markedly lower voxels-per-RESEL estimates across all qmaps and tissues, reflecting a reduced effective smoothness and a more spatially fragmented statistical topology.

Differences between parametric inference under stationary and non-stationary assumptions remained minimal for all smoothing strategies. Consistent with the very similar effective smoothness estimated under both frameworks, the corresponding RESEL estimates also differed only marginally. Non-parametric inference did not provide voxels-per-RESEL estimates, resulting in missing values for all smoothing approaches, as RESEL-based smoothness estimation is not part of the SnPM inference framework.

The number of significant voxels and clusters per GLM and inference approaches, for all smoothed qMRI maps are summarized in Table 4. Across all inference strategies, TWS consistently yielded the highest number of statistically significant voxels (**nSigVox**) across quantitative MRI parameters and tissue classes, whereas gTSPOON generally produced intermediate values and SUSANs systematically resulted in the lowest extent of significant activation. This trend was particularly pronounced for MTsat and R2star maps, where TWS identified substantially larger significant regions than SUSANs in both GM and WM.

**Table 4:**
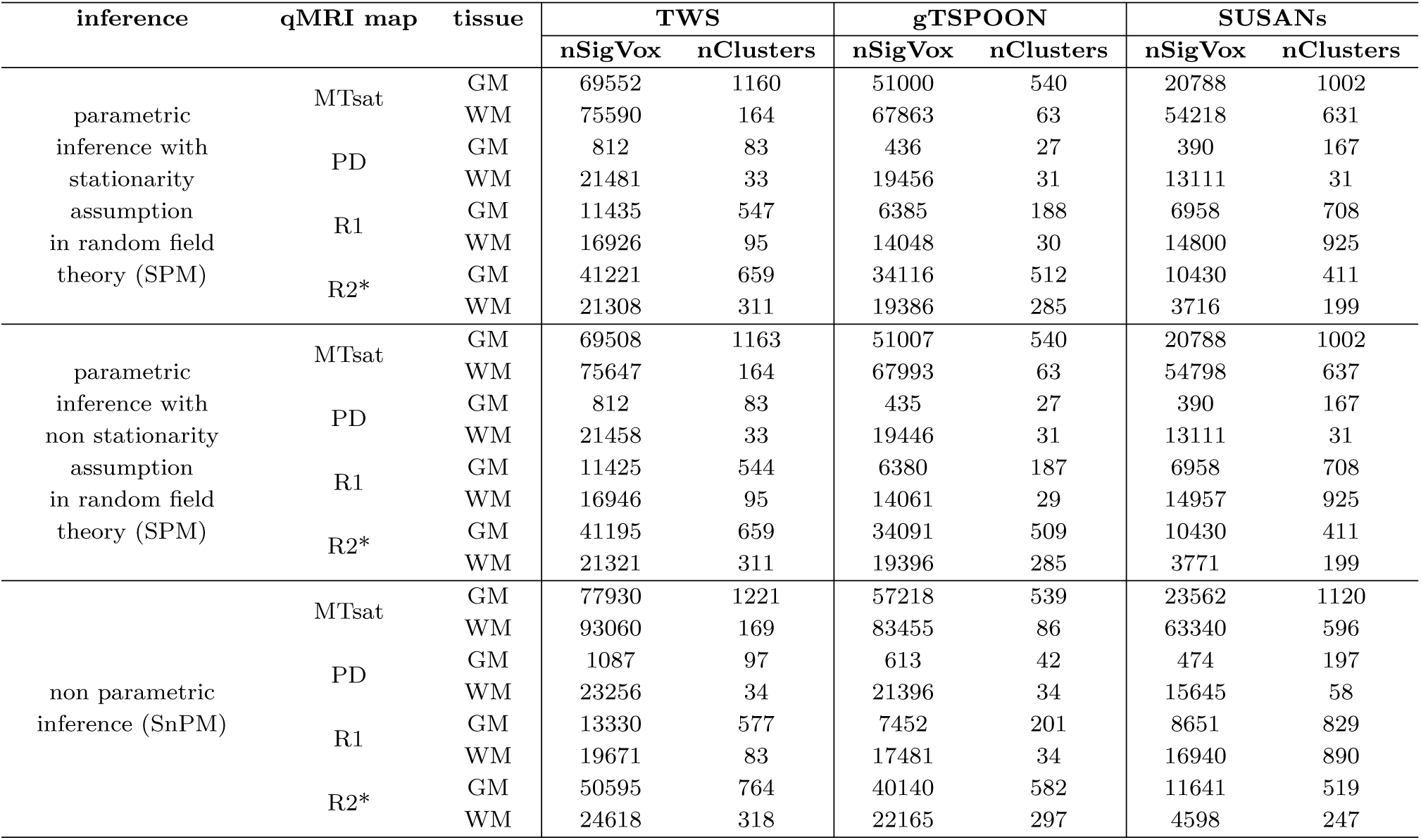
Number of statistically significant voxels and clusters (*p <* 0.05 FWER) for TWS, gTSPOON and SUSANs for each smoothing approach across inference’s approaches, quantitative parameters and tissue types.

For MTsat maps, the largest effects were observed in WM, with TWS reaching up to 93,060 significant voxels under non-parametric inference, compared with 83,455 for gTSPOON and 63,340 for SUSANs. Similarly, in GM, TWS consistently produced the largest significant extent, followed by gTSPOON and then SUSANs. Despite these differences in voxel counts, SUSANs often generated a comparatively large number of clusters, particularly in GM (e.g., 1,120 clusters for MTsat under SnPM), indicating a more spatially fragmented statistical topology.

PD map analyses showed overall lower numbers of significant voxels, especially in GM, where all smoothing approaches produced relatively sparse detections. Nevertheless, TWS remained associated with the highest sensitivity, while SUSANs generally produced the largest cluster fragmentation. In WM, PDmap results exhibited markedly larger significant extents than GM across all methods and inference approaches.

For R1 map, TWS again produced the largest significant voxel counts across tissues and inference methods, whereas SUSANs exhibited notably elevated cluster numbers despite lower voxel extents. This pattern was especially evident in WM, where SUSANs generated up to 925 clusters under parametric inference despite substantially fewer significant voxels than TWS, suggesting increased spatial dispersion and fragmentation of detected regions.

R2* maps demonstrated strong differences between smoothing strategies. TWS and gTSPOON yielded relatively similar voxel counts, whereas SUSANs consistently showed a marked reduction in significant voxels, particularly in WM (e.g., 3,716 significant voxels for SUSANs versus 21,308 for TWS under stationary parametric inference). However, SUSANs still produced a relatively high number of clusters, again indicating a fragmented spatial organization of significant effects.

Across inference frameworks, results obtained using parametric inference with and without stationarity assumptions were nearly identical, with only negligible variations in significant voxel and cluster counts. In contrast, non-parametric inference (SnPM) systematically increased the number of significant voxels and clusters across all smoothing approaches, tissues, and qMRI maps, reflecting the generally higher sensitivity of permutation-based inference in this dataset.

### 3.2 Effect of smoothing on continuous SPMs

The smoothing strategy substantially influences the statistical properties of quantitative MRI analyses, both at the voxel-wise model-fitting level and at the level of statistical inference.

#### 3.2.1 Voxel-wise log-likelihood comparison

Since the voxel-wise log-likelihood (LL) is computed from the GLM’s parameter and residual maps, it does not depend on the inference scheme. Thus, the log-likelihood maps make it possible to highlight the minimization of residuals during GLM estimation as a function of the different smoothing approaches, see Figure 3. The TWS-smoothed MTsat maps lead to the largest LL values mostly in GM or in the core of WM; while gTSPOON-smoothed MTsat maps have the the largest LL at in the more superficial voxels of the GM and WM masks. Then regions in which SUSANs-smoothed MTsat maps have a better fit are located at the very boundaries between GM and WM.

**Figure 3:**
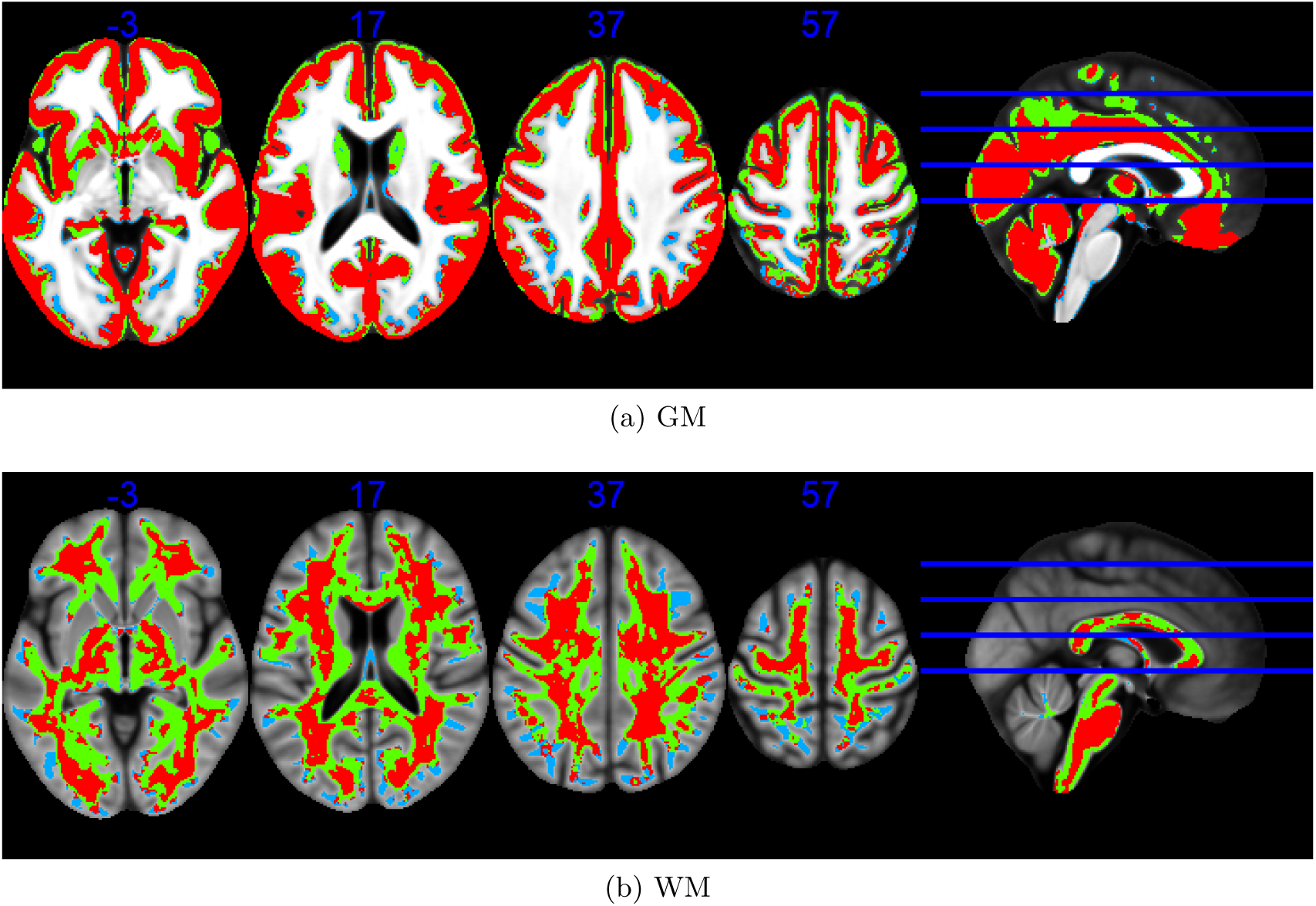
Voxelwise log-likelyhood maps, in GM (top) and WM (bottom) identifying regions in which GLM’s residuals are minimized on the MTsat maps smoothed using TWS (red), gTSPOON (green), and SUSANs (blue). The four axial slices are located at z =-3, 17, 37 and 57 mm, from left to right, as illustrated on the sagital slice (right).

Bland-Altman plots are computed between the voxel-wise log-likelihood maps of each smoothing approach for MTsat quantitative parameter, see Figure 4. To summarize the behavior of the Bland–Altman plots for the other quantitative parameters, the 95% limits of agreement (LoA) are fully determined by the mean difference (*d*) and the standard deviation of the differences (*SD_d_*), according to: *LoA*_95%_ = *d* ± 1.96*, SD_d_*. Therefore, Bland–Altman plots can be qualitatively interpreted through the corresponding values of *d* and *SD_d_*. Table 5 summarizes these mean differences and standard deviations across all quantitative parameters.

**Figure 4:**
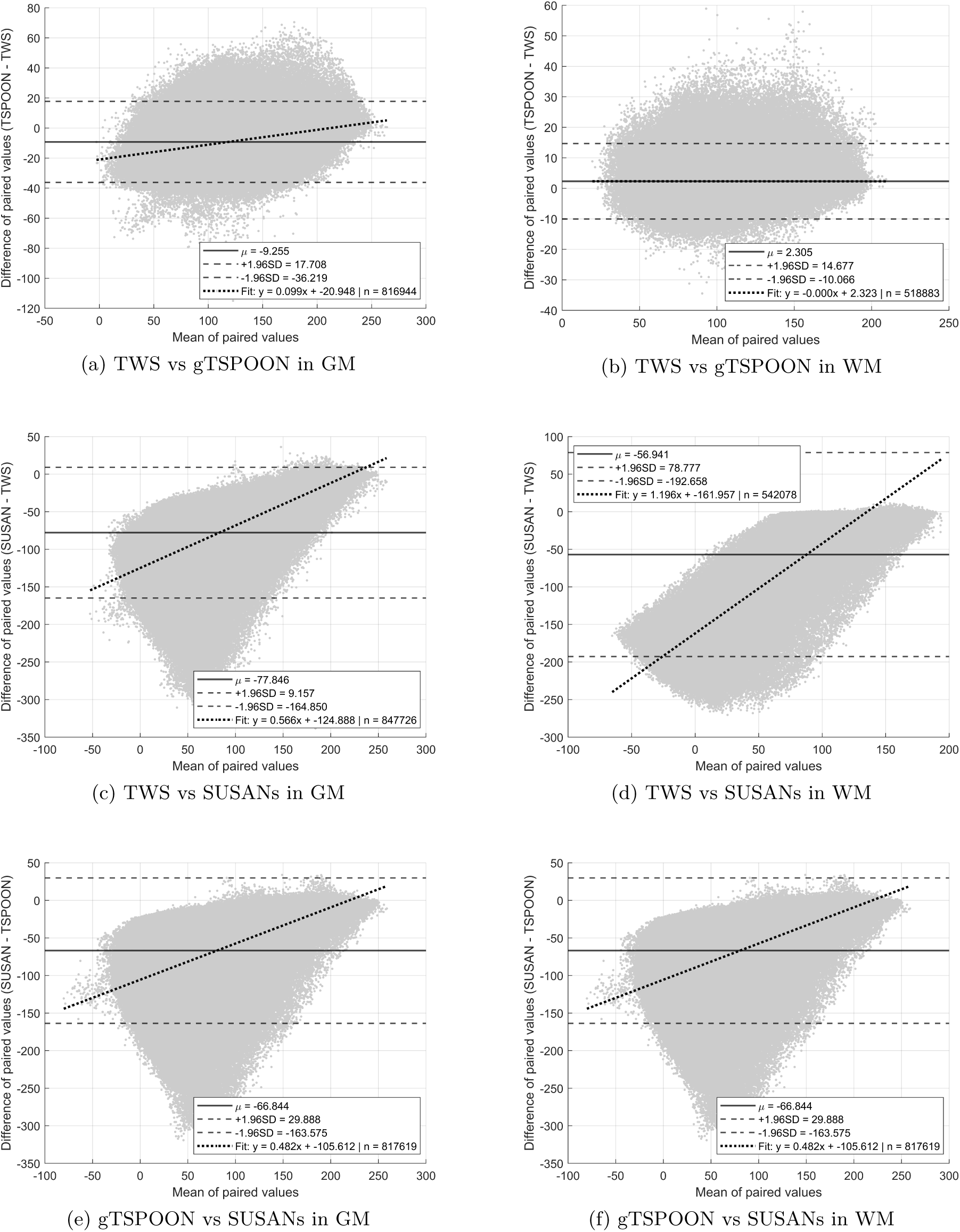
Bland-Altman plots of GLM estimation MTsat maps.

**Table 5:** Overview of mean differences (*d*) and standard deviations of the differences (*SD_d_*) computed from Bland–Altman analyses of voxelwise GLM log-likelihood maps across smoothing approaches, quantitative MRI parameters, and tissue classes.

| agreement | qmap | tissue | d | $SD_d$ |
| --- | --- | --- | --- | --- |
| TWS<br>vs<br>gTSPOON | MTsat | GM | -9.255 | 13.757 |
|  |  | WM | 2.305 | 6.312 |
|  | PD | GM | -7.042 | 9.179 |
|  |  | WM | -1.941 | 6.178 |
|  | R1 | GM | -8.192 | 11.709 |
|  |  | WM | 0.625 | 4.583 |
|  | R2* | GM | -5.672 | 8.421 |
|  |  | WM | -0.136 | 2.666 |
| TWS<br>vs<br>SUSANs | MTsat | GM | -77.846 | 44.389 |
|  |  | WM | -56.941 | 69.244 |
|  | PD | GM | -136.539 | 121.602 |
|  |  | WM | -99.667 | 134.371 |
|  | R1 | GM | -86.877 | 68.501 |
|  |  | WM | -64.359 | 83.576 |
|  | R2* | GM | -73.012 | 55.634 |
|  |  | WM | -50.361 | 69.362 |
| gTSPOON<br>vs<br>SUSANs | MTsat | GM | -66.844 | 49.353 |
|  |  | WM | -53.719 | 68.681 |
|  | PD | GM | -123.639 | 121.100 |
|  |  | WM | -88.261 | 130.350 |
|  | R1 | GM | -75.367 | 71.851 |
|  |  | WM | -58.382 | 80.629 |
|  | R2* | GM | -64.703 | 53.588 |
|  |  | WM | -44.934 | 64.868 |

Across all quantitative MRI parameters, the highest agreement was consistently observed between TWS and gTSPOON, as indicated by comparatively small mean differences (*d*) and reduced standard deviations of the differences (*SD_d_*). In this comparison, WM systematically exhibited lower variability than GM, with particularly small dispersions for R2* (*SD_d_* = 2.666) and R1 (*SD_d_* = 4.583), suggesting a strong consistency of voxel-wise GLM log-likelihood distributions between these two smoothing strategies.

In contrast, comparisons involving SUSANs demonstrated substantially larger negative biases and markedly wider dispersions across all qMRI maps and tissue classes. The strongest discrepancies were observed for PD maps, where TWS vs SUSANs reached mean differences of *d* = −136.539 in GM and *d* = −99.667 in WM, associated with very large variability (*SD_d_* = 121.602 and 134.371, respectively). Similar trends were observed for gTSPOON vs SUSANs, although the magnitude of the bias was generally slightly reduced compared with TWS vs SUSANs.

Across all agreements, GM tended to show larger absolute mean differences than WM, indicating a larger sensitivity of GM voxel-wise statistics to the choice of smoothing approach. Likewise, variability was generally elevated in comparisons involving SUSANs, particularly for PD and R1 maps, whereas R2* consistently showed the lowest dispersion among the quantitative parameters. Overall, these results indicate that TWS and gTSPOON produce highly comparable voxel-wise GLM log-likelihood maps, while SUSANs introduces substantially different statistical distributions, especially in GM regions.

#### 3.2.2 Voxel-wise T-value comparison

The statistical T-value maps are generated prior to statistical inference and therefore do not depend on whether inference is parameterized or not. This makes it possible to compare the impact of different smoothing approaches on the statistical analysis independently of the inference step. Figure 5 shows the Bland–Altman plots computed on the statistical T-values maps for MTsat quantitative maps. As for the log-likelihood maps comaprison, the Bland–Altman plots are qualitatively interpreted through the the mean differences *d* and standard deviations *SD_d_* across all quantitative parameters, as summarized in Table 6.

**Figure 5:**
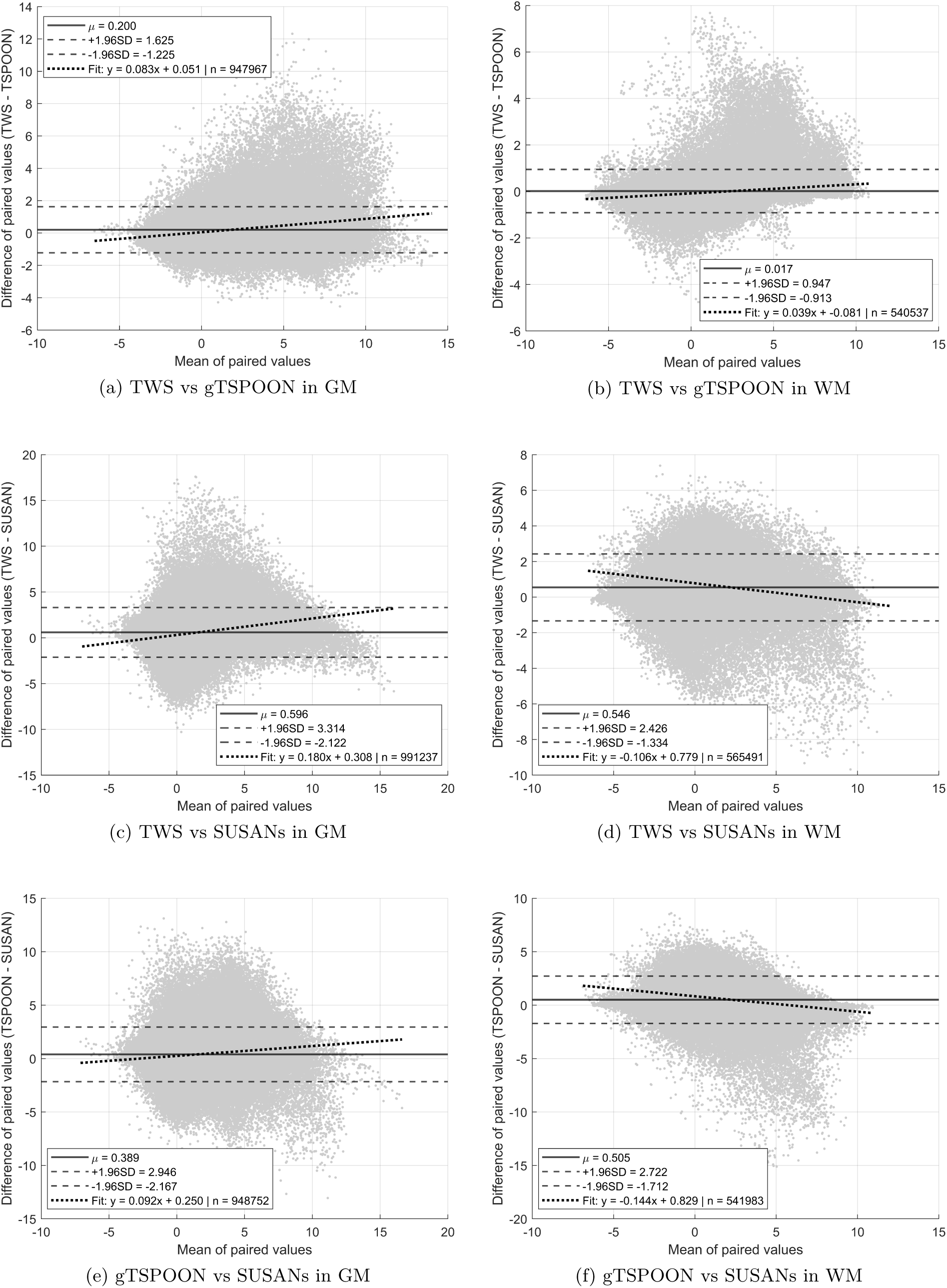
Bland-Altman plots of the statistical T-values maps for MTsat quantitative parameter.

**Table 6:** Overview of difference mean (*d*) and standard deviation (*SD_d_*) computed in Bland-Altman plots obtained from T-values maps for the parametric or non-parametric statistical inference approaches (PI or non-PI) and with or without stationarity assumption (SA or non-SA) in RFT, across smoothing approaches, qMRI maps, and tissue types.

| agreement | qMRI map | tissue | PI with SA in RFT |  | PI with non-SA in RFT |  | non-PI |  |
| --- | --- | --- | --- | --- | --- | --- | --- | --- |
| | | | $d$ | $SD_d$ | $d$ | $SD_d$ | $d$ | $SD_d$ |
| TWS<br>vs<br>gTSPOON | MTsat | GM | 0.200 | 0.727 | 0.200 | 0.727 | 0.200 | 0.727 |
|  |  | WM | 0.017 | 0.475 | 0.017 | 0.475 | 0.017 | 0.475 |
|  | PD | GM | -0.110 | 0.607 | -0.110 | 0.607 | -0.110 | 0.607 |
|  |  | WM | 0.083 | 0.379 | 0.083 | 0.379 | 0.083 | 0.379 |
|  | R1 | GM | 0.047 | 0.640 | 0.047 | 0.640 | 0.047 | 0.640 |
|  |  | WM | 0.060 | 0.368 | 0.060 | 0.368 | 0.060 | 0.368 |
|  | R2* | GM | 0.175 | 0.542 | 0.175 | 0.542 | 0.175 | 0.542 |
|  |  | WM | -0.035 | 0.170 | -0.035 | 0.170 | -0.035 | 0.170 |
| TWS<br>vs<br>SUSANs | MTsat | GM | 0.596 | 1.387 | 0.596 | 1.387 | 0.596 | 1.387 |
|  |  | WM | 0.546 | 0.959 | 0.546 | 0.959 | 0.546 | 0.959 |
|  | PD | GM | -0.330 | 1.886 | -0.330 | 1.886 | -0.330 | 1.886 |
|  |  | WM | -0.293 | 1.974 | -0.293 | 1.974 | -0.293 | 1.974 |
|  | R1 | GM | -0.328 | 1.324 | -0.328 | 1.324 | -0.328 | 1.324 |
|  |  | WM | 0.111 | 0.832 | 0.111 | 0.832 | 0.111 | 0.832 |
|  | R2* | GM | 1.155 | 1.456 | 1.155 | 1.456 | 1.155 | 1.456 |
|  |  | WM | 0.548 | 1.224 | 0.548 | 1.224 | 0.548 | 1.224 |
| gTSPOON<br>vs<br>SUSANs | MTsat | GM | 0.389 | 1.304 | 0.389 | 1.304 | 0.389 | 1.304 |
|  |  | WM | 0.505 | 1.131 | 0.505 | 1.131 | 0.505 | 1.131 |
|  | PD | GM | -0.217 | 1.666 | -0.217 | 1.666 | -0.217 | 1.666 |
|  |  | WM | -0.345 | 1.744 | -0.345 | 1.744 | -0.345 | 1.744 |
|  | R1 | GM | -0.376 | 1.282 | -0.376 | 1.282 | -0.376 | 1.282 |
|  |  | WM | 0.056 | 0.979 | 0.056 | 0.979 | 0.056 | 0.979 |
|  | R2* | GM | 0.976 | 1.457 | 0.976 | 1.457 | 0.976 | 1.457 |
|  |  | WM | 0.516 | 1.182 | 0.516 | 1.182 | 0.516 | 1.182 |

The Bland–Altman analysis revealed identical agreement metrics across all statistical inference approaches, including parametric inference with or without the stationarity assumption in RFT and non-parametric inference. This indicates that the inference strategy did not affect the agreement between statistical T-value maps derived from the different smoothing approaches.

Across all quantitative maps, the strongest agreement was consistently observed between TWS and gTSPOON, as reflected by lower mean differences (*d*) and smaller standard deviations (*SD_d_*). In contrast, comparisons involving SUSANs exhibited larger biases and wider dispersions, particularly within GM of R2* quantitative maps. Overall, WM generally showed lower variability than GM, suggesting a higher consistency of statistical T-value distributions across smoothing approaches in WM.

#### 3.2.3 Voxel-wise thresholded T-maps comparison

The agreement between the different smoothing approaches can also be assessed using the Jaccard, Dice, and Cohen indices. These similarity metrics were computed between the statistically significant regions obtained from the GLM applied to the differently smoothed maps. Since these metrics were calculated after statistical inference, they were evaluated separately for each of the three inference approaches, see Table 7.

**Table 7:** Agreement metrics (Jaccard, Dice, and Cohen’s *κ*) between statistically significant regions (*p <* 0.05 FWE) for parametric or non-parametric statistical inference approaches (PI or non-PI) and with or without stationarity assumption (SA or non-SA) in RFT, across smoothing approaches, qMRI maps, and tissue types.

| agreement | qMRI map | tissue | PI with SA in RFT |  |  | PI with non-SA in RFT |  |  | non-PI |  |  |
| --- | --- | --- | --- | --- | --- | --- | --- | --- | --- | --- | --- |
| | | | Jaccard | Dice | $\kappa$ | Jaccard | Dice | $\kappa$ | Jaccard | Dice | $\kappa$ |
| TWS<br>vs<br>gTSPOON | MTsat | GM | 0.600 | 0.750 | 0.748 | 0.600 | 0.750 | 0.748 | 0.610 | 0.758 | 0.756 |
|  |  | WM | 0.882 | 0.937 | 0.937 | 0.883 | 0.938 | 0.937 | 0.885 | 0.939 | 0.938 |
|  | PD | GM | 0.296 | 0.457 | 0.457 | 0.296 | 0.457 | 0.457 | 0.290 | 0.449 | 0.449 |
|  |  | WM | 0.893 | 0.944 | 0.944 | 0.894 | 0.944 | 0.944 | 0.898 | 0.946 | 0.946 |
|  | R1 | GM | 0.403 | 0.575 | 0.574 | 0.404 | 0.575 | 0.575 | 0.415 | 0.587 | 0.586 |
|  |  | WM | 0.813 | 0.897 | 0.896 | 0.812 | 0.896 | 0.896 | 0.814 | 0.897 | 0.897 |
|  | R2* | GM | 0.731 | 0.845 | 0.844 | 0.731 | 0.845 | 0.844 | 0.712 | 0.831 | 0.830 |
|  |  | WM | 0.756 | 0.861 | 0.861 | 0.756 | 0.861 | 0.860 | 0.762 | 0.865 | 0.865 |
| TWS<br>vs<br>SUSANs | MTsat | GM | 0.276 | 0.432 | 0.430 | 0.276 | 0.432 | 0.430 | 0.280 | 0.437 | 0.434 |
|  |  | WM | 0.682 | 0.811 | 0.809 | 0.689 | 0.816 | 0.814 | 0.655 | 0.791 | 0.789 |
|  | PD | GM | 0.069 | 0.130 | 0.130 | 0.069 | 0.130 | 0.130 | 0.066 | 0.124 | 0.124 |
|  |  | WM | 0.608 | 0.756 | 0.756 | 0.609 | 0.757 | 0.756 | 0.668 | 0.801 | 0.800 |
|  | R1 | GM | 0.299 | 0.460 | 0.459 | 0.299 | 0.460 | 0.459 | 0.310 | 0.473 | 0.472 |
|  |  | WM | 0.619 | 0.765 | 0.764 | 0.624 | 0.769 | 0.768 | 0.632 | 0.774 | 0.774 |
|  | R2* | GM | 0.242 | 0.390 | 0.389 | 0.242 | 0.390 | 0.389 | 0.220 | 0.361 | 0.359 |
|  |  | WM | 0.168 | 0.288 | 0.287 | 0.170 | 0.291 | 0.290 | 0.178 | 0.303 | 0.302 |
| gTSPOON<br>vs<br>SUSANs | MTsat | GM | 0.277 | 0.434 | 0.432 | 0.277 | 0.434 | 0.431 | 0.285 | 0.444 | 0.441 |
|  |  | WM | 0.665 | 0.799 | 0.797 | 0.671 | 0.803 | 0.801 | 0.652 | 0.789 | 0.787 |
|  | PD | GM | 0.100 | 0.182 | 0.182 | 0.100 | 0.182 | 0.182 | 0.094 | 0.171 | 0.171 |
|  |  | WM | 0.671 | 0.803 | 0.803 | 0.671 | 0.803 | 0.803 | 0.725 | 0.841 | 0.841 |
|  | R1 | GM | 0.243 | 0.391 | 0.390 | 0.243 | 0.391 | 0.391 | 0.254 | 0.405 | 0.404 |
|  |  | WM | 0.556 | 0.715 | 0.714 | 0.559 | 0.717 | 0.717 | 0.545 | 0.706 | 0.705 |
|  | R2* | GM | 0.294 | 0.454 | 0.453 | 0.294 | 0.455 | 0.453 | 0.277 | 0.434 | 0.432 |
|  |  | WM | 0.185 | 0.313 | 0.312 | 0.188 | 0.316 | 0.315 | 0.198 | 0.330 | 0.329 |

Across all pairwise comparisons between smoothing approaches, the similarity measures are highly consistent across parametric inference, with or without the stationarity assumption in RFT, as well as with the non-parametrics. Specifically, the differences observed between the two stationarity conditions are on the order of at most (10^−3^). In contrast, the differences in similarity measures between parametric and non-parametric inference reach an order of magnitude of up to.01. These findings indicate that statistically significant regions (*p <* 0.05 FWE-corrected) are more consistent when varying the stationarity assumption within RFT than when comparing parametric versus non-parametric inference approaches. Nevertheless, the overall differences in similarity measures remain relatively small, suggesting a general consistency in the spatial overlap of statistically significant regions across the different inference methods.

Regarding the smoothing approaches, statistically significant regions exhibited relatively higher similarity between TWS and gTSPOON, particularly within white matter (WM). However, these similarity values remained moderate, especially in gray matter (GM), indicating that the agreement between smoothing approaches was not uniformly high across the brain. Conversely, similarity measures were further reduced when comparing either TWS or gTSPOON with SUSANs, particularly in GM regions. Overall, similarity measures were consistently higher in WM than in GM across all quantitative parameters, with the exception of R2*, for which comparisons between TWS/gTSPOON and SUSANs showed the opposite trend.

## 4 Discussion

This study investigated the impact of tissue-specific smoothing strategies on voxel-wise quantitative MRI analyses by comparing TWS, gTSPOON, and SUSANs across multiple quantitative parameters, tissue classes, and statistical inference frameworks.

### 4.1 Smoothing effects across inference approaches, qMRI maps and tissue classes

Reproducing the age-related findings of Callaghan et al. [6] using three different tissue-specific smoothing approaches revealed systematic and interpretable differences in statistical outcomes depending on the chosen smoothing approach. TWS, from the original paper, and gTSPOON produced broadly comparable spatial patterns of age-related variations depending on the quantitative parameters, consistent with the original study[6]. Yet TWS yielded a notably higher number of significant voxels and clusters (Table 4). This difference can be attributed to the slightly higher effective smoothing achieved with TWS, which reduces the number of RESELs (Table 3) and also the multiple comparison problem, thus leading to slightly increased sensitivty (Table 2). The similarity between TWS and gTSPOON suggests that both linear and nonlinear tissue-weighting modulation preserve the essential spatial features of the underlying signal when the effective smoothness is comparable.

By contrast, SUSANs produced results that differed markedly from the other two approaches. SUSANs showed substantially fewer significant voxels and more significant clusters (Table 4), which reflects its considerably lower effective smoothness, approximately half that of TWS and gTSPOON. This reduced smoothing led to a smaller number of voxels per RESEL (Table 3) and, consequently, a more conservative statistical outcome (Table 2). Unlike TWS and gTSPOON, SUSANs includes intensity modulation (Eq. 5). This modulation preserves local contrast and sharp transitions between gray and white matter, which helps to retain anatomical detail that conventional smoothing tends to blur.

The present results demonstrate that the choice of smoothing strategy substantially influences the statistical properties of quantitative MRI analyses, both at the voxel-wise model-fitting level and at the level of statistical inference. Across all analyses, the highest agreement was consistently observed between TWS and gTSPOON, whereas SUSANs produced markedly different statistical distributions and spatial patterns, particularly within gray matter.

At the voxel-wise level, the comparison of GLM log-likelihood maps revealed distinct spatial behaviors associated with each smoothing strategy. The voxel-wise log-likelihood maps (Figs. 3 19, 20, 21) further support this interpretation. The regions where TWS-smoothed maps yield the best GLM fit are mainly located within GM, whereas the regions where SUSAN-smoothed maps perform best lie at the GM–WM boundaries, particularly within sulcal and gyral transitions. In WM, the pattern varies across quantitative parameters. On the PD maps, TWS predominates in WM. In contrast, the R2* maps show a predominance of regions where gTSPOON-smoothed maps provide the best GLM fit. The R1 maps exhibit an intermediate behavior between TWS and gTSPOON within WM.

On all the smoothed maps (i.e. MTsat, PD, R1 and R2*), it can also be observed that TWS tends to perform best in the central portions of tissues, whether GM or WM, while gTSPOON tends to perform better toward the periphery of tissue regions, excluding the tissue boundaries where SUSAN dominates. On the one hand, this suggests that the intensity-modulated smoothing of SUSANs preserves local signal intensity gradients, enabling the model to capture sharp tissue transitions and large variations in quantitative values more accurately. On the other hand, the continuous tissue weighting in TWS may better capture subtle variations within tissue, whereas the binary tissue mask used in gTSPOON is more suited to the homogeneity of the tissue.

Bland–Altman analyses (Table 5) demonstrated close agreement between the two tissue-weighted smoothing approaches (TWS and gTSPOON), with mean differences close to zero and relatively small standard deviations across all quantitative MRI parameters. In contrast, comparisons involving SUSANs showed substantially larger negative mean differences, indicating that SUSANs consistently produced higher voxelwise GLM log-likelihoods than either tissue-weighted method. These differences were generally larger in gray matter than white matter and were most pronounced for PD maps, suggesting that PD-derived likelihood estimates are more sensitive to the choice of smoothing approach. The larger standard deviations observed for comparisons involving SUSANs further indicate greater spatial variability in the differences between smoothing methods.

The analyses performed on T-statistics maps pointed at similar conclusions. The agreement between smoothing approaches remained largely independent of the inference strategy, as identical Bland–Altman metrics were obtained for parametric inference with or without stationarity assumptions and for non-parametric inference. This indicates that the primary source of variability originates from the smoothing step itself rather than from the subsequent statistical inference procedure. Such findings emphasize the critical influence of preprocessing choices on first-level statistical estimates in quantitative MRI analyses.

Regarding the thresholded statistical maps, the similarity analyses based on Jaccard, Dice, and Cohen indices demonstrated a high overall consistency of statistically significant regions across inference methods. The small differences observed between stationary and non-stationary RFT corrections suggest that relaxing the stationarity assumption has only a limited impact on the final spatial distribution of significant effects in the present dataset. Slightly larger discrepancies were observed between parametric and non-parametric inference, pointing at a slightly increased sensitivity for the non-parametric approach, although these differences remained relatively modest overall. Together, these observations support the robustness of the reported statistical findings across inference frameworks.

With respect to the smoothing approaches, the thresholded maps again showed stronger overlap between TWS and gTSPOON, particularly in WM regions. The reduced similarity observed when comparing either tissue-weighted approach with SUSANs further supports the hypothesis that tissue-informed smoothing strategies preserve comparable anatomical and statistical structures, whereas SUSANs produces systematically different spatial distributions of significant effects. The generally higher overlap observed in WM compared with GM additionally highlights the greater stability of smoothing procedures within structurally homogeneous regions. The exception observed for R2* may reflect the increased sensitivity of this parameter to local susceptibility variations and iron-related microstructural heterogeneity, which can alter the spatial behavior of smoothing differently across tissue classes.

Overall, these findings indicate that tissue-specific weighted smoothing approaches produce more consistent statistical behavior than conventional edge-preserving smoothing methods in quantitative MRI analyses. Still in the absence of ground truth for the observed effect, the strong agreement observed between TWS and gTSPOON suggests that incorporating tissue priors into the smoothing process contributes to stabilizing voxel-wise model estimation and preserving the spatial consistency of statistical effects. In contrast, the larger discrepancies associated with SUSANs indicate that smoothing strategies not explicitly constrained by tissue information may substantially affect statistical distributions, particularly in anatomically heterogeneous cortical regions. These results underline the importance of carefully selecting smoothing strategies in quantitative MRI studies, as preprocessing choices may directly influence both voxel-wise statistical estimates and the spatial extent of inferred effects.

### 4.2 Inference parametrization across smoothing approaches, quantitative parameters and tissue classes

Although the present study was not primarily designed to compare statistical inference frameworks, the comparison between parametric and non-parametric inference frameworks revealed an overall consistency of statistical results across all quantitative MRI parameters, tissue classes, and smoothing strategies. Parametric inference performed under stationary and non-stationary random field theory (RFT) assumptions produced nearly identical statistical thresholds, RESEL estimations, and spatial distributions of significant effects (Tables 2 and 3). The very small differences observed between these two RFT implementations indicate that violations of the stationarity assumption had only a minimal impact on statistical inference in the present dataset. This observation further suggests that the effective spatial smoothness introduced by the different smoothing strategies remained sufficiently stable across the brain to ensure robust RFT-based correction.

In contrast, non-parametric inference using SnPM systematically yielded lower corrected T-thresholds and larger extents of statistically significant voxels compared with parametric inference (Tables 2, **??**, and 4). These findings suggest a moderately increased sensitivity of permutation-based inference relative to RFT approaches. Nevertheless, despite these quantitative differences in statistical sensitivity, the spatial organization of significant effects remained highly consistent across inference frameworks. Similarity analyses based on Jaccard, Dice, and Cohen indices demonstrated only modest discrepancies between parametric and non-parametric inference, with differences remaining on the order of 10^−2^ (Table 1). Likewise, Bland–Altman analyses of statistical T-value maps (Figure 5) showed identical agreement metrics independently of the inference framework, indicating that the primary source of variability originated from the smoothing procedure itself rather than from the subsequent statistical inference method.

Importantly, the relative behavior of the smoothing strategies remained stable across all inference frameworks. TWS and gTSPOON consistently exhibited the strongest agreement and the largest overlap of significant regions, whereas SUSANs systematically produced more fragmented statistical topologies characterized by fewer significant voxels and a larger number of clusters (Table 4). These observations indicate that preprocessing choices, particularly the effective spatial smoothness induced by tissue-specific smoothing strategies, exert a substantially greater influence on voxel-wise statistical outcomes than the choice between parametric and non-parametric inference frameworks.

### 4.3 Reproducibility considerations for SUSANs

The present implementation of SUSANs was designed with particular attention to reproducibility considerations highlighted in previous work. Chen et al. [9] demonstrated that subtle implementation differences in the parameterization of SUSAN smoothing between FSL and Nipype can lead to substantial discrepancies in smoothed images and downstream statistical analyses. In particular, very small numerical variations in the spatial smoothing parameter *d_t_* were shown to propagate non-linearly through the exponential weighting function of SUSAN, resulting in impactful different voxel contributions within local neighbourhoods.

In line with Chen et al. [9], our implementation avoids wrapper-induced variability by using the FSL command-line binary directly, thus eliminating discrepancies between FSL and Nipype parameterizations. Our parameterization is further justified because it places SUSANs on the same effective smoothing scale as the other methods, with a target FWHM of 3 mm converted into *d_t_* ≈ 1.27 mm. Doing so improves methodological comparability and helps reduce inter-method variability. In addition, estimating the brightness threshold from valid voxel intensities provides a data-driven and modality-specific setting that is less sensitive to artefactual values. This strengthens the robustness of SUSANs within a qMRI pipeline, as the smoothing then depends on the actual statistical properties of the data.

## 5 Conclusion

Overall, TWS and gTSPOON demonstrated the highest consistency in voxel-wise model fitting, statistical topology, and spatial overlap of significant effects, supporting the benefit of incorporating tissue information during smoothing. In contrast, SUSANs produced distinct statistical patterns, but voxel-wise log-likelihood analyses suggest that these differences reflect preferential performance in specific anatomical configurations rather than a systematic limitation of intensity-based smoothing.

A major contribution of this work is therefore the introduction of voxel-wise GLM log-likelihood mapping as a framework for spatially characterizing smoothing performance. Unlike conventional global comparisons, voxel-wise LL maps revealed that each smoothing approach possesses specific “preferred regions” where it provides the best local model fit.

This suggest that no single smoothing strategy is uniformly optimal across the entire brain. Instead, the optimal preprocessing approach appears to depend on local anatomical organization, tissue heterogeneity, and the underlying biological process being investigated. In the specific context of age-related qMRI analyses, this finding is particularly relevant because age effects are known to vary spatially across tissue compartments and cortical structures. Consequently, voxel-wise LL mapping provides a principled framework for guiding the selection of smoothing strategies according to the neuroanatomical regions or tissue classes of interest, thereby maximizing statistical sensitivity while preserving anatomical specificity.

Importantly, the present results also demonstrate that the influence of smoothing extends beyond simple changes in effective smoothness or statistical thresholds. The observed regional preferences indicate that smoothing strategies modify the local balance between noise reduction, tissue preservation, and edge conservation in fundamentally different ways. This effect was consistently observed across all quantitative parameters and remained stable across both parametric and non-parametric inference frameworks, emphasizing that preprocessing choices themselves constitute a major determinant of the resulting statistical topology.

More broadly, the proposed voxel-wise LL framework may serve as a general tool for evaluating and optimizing preprocessing pipelines in quantitative neuroimaging. Rather than relying on a single globally applied smoothing kernel, future work could investigate adaptive or hybrid smoothing strategies in which the locally optimal smoothing approach is selected voxel-wise according to model likelihood. Such adaptive frameworks could combine the complementary strengths of tissue-specific and edge-preserving methods, potentially improving both anatomical precision and statistical sensitivity in quantitative MRI studies.

## Acknowledgements, Author Contributions, Funding and Competing Interests

## 5.1 Acknowledgements

The authors would like to thank Martina F. Callaghan for sharing the data prior to the open-access release and for her contributions to data curation and resource management. The authors also thank Thomas E. Nichols for his expertise in statistical inference and for initiating the integration of SUSANs into this research methodology.

## 5.2 Author Contributions

AJ: Conceptualization, Methodology, Software, Validation, Formal analysis, Investigation, Writing the original draft, Visualization

CP: Conceptualization, Methodology, Resources, Writing (review & editing), Supervision, Project administration

## 5.3 Funding and Competing Interests

Both AJ and CP are supported by the F.R.S.-FNRS, Belgium. No external funding was received for this study. The authors declare that they have no conflict of interest.

## A Supplementary Material

### A.1 Smoothing effects across different aging model parameterizations

To further investigate the impact of smoothing approaches on statistical parametric maps in the context of non-linear aging effects, a quadratic (centered) age term was added to the GLM. Statistical inference was performed using an F-test under the assumption of non-stationarity in RFT. Statistically significant non-linear effects of age are shown in Figure 6 for the MTsat qMRI maps, while Tables 8 and 9 report the statistical T-thresholds (*p <* 0.05 FWE) and the mean number of voxels per RESEL for all qMRI maps and tissue types.

**Figure 6:**
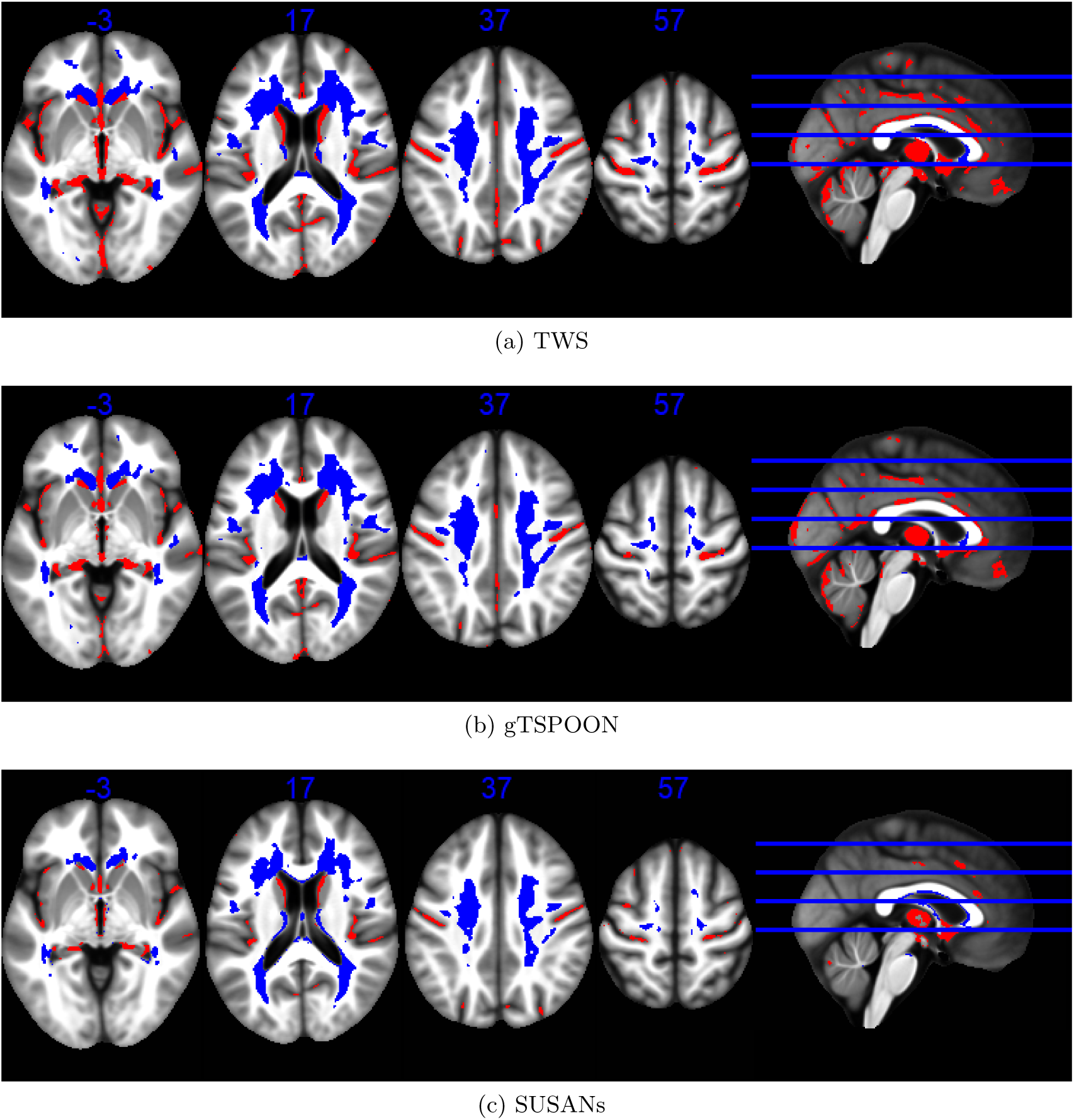
Statistical parametric maps identifying regions in GM (red) and WM (blue) in which MTsat significantly varied with age at the *p <* 0.05 FWE corrected level using non parametric statistical inference (SnPM). The results are superimposed on the mean MT map for the cohort in MNI space. The four axial slices are located at z = - 3, 17, 37 and 57 mm, from left to right, as illustrated on the sagital slice (right).

**Table 8:** Overview of statistical F-threshold values after parametric inference under non-stationarity hypothesis (SPM) (*p <* 0.05 FWER) for any age effect, including the quadratic age term, across different smoothing approaches, quantitative parameters, and tissue types.

| Smoothing | MTsat |  | PD |  | R1 |  | R2* |  |
| --- | --- | --- | --- | --- | --- | --- | --- | --- |
|  | GM | WM | GM | WM | GM | WM | GM | WM |
| TWS | 17.78 | 15.78 | 17.20 | 16.46 | 17.48 | 15.38 | 17.44 | 15.60 |
| gTSPOON | 17.91 | 15.91 | 17.29 | 16.50 | 17.58 | 15.46 | 17.46 | 15.55 |
| SUSANs | 19.17 | 17.81 | 19.17 | 18.50 | 19.17 | 17.88 | 19.17 | 18.30 |

**Table 9:** Overview of number of voxels per RESEL after parametric inference under non-stationarity hypothesis (SPM) (*p <* 0.05 FWER) for any age effect, including the quadratic age term, across different smoothing approaches, quantitative parameters, and tissue types.

| Smoothing | MTsat |  | PD |  | R1 |  | R2* |  |
| --- | --- | --- | --- | --- | --- | --- | --- | --- |
|  | GM | WM | GM | WM | GM | WM | GM | WM |
| TWS | 171.89 | 447.08 | 270.15 | 258.53 | 218.09 | 621.89 | 224.89 | 516.99 |
| gTSPOON | 148.99 | 379.77 | 241.18 | 238.49 | 545.54 | 545.54 | 210.38 | 506.86 |
| SUSANs | 21.16 | 98.14 | 29.64 | 40.18 | 27.14 | 93.69 | 21.03 | 67.90 |

The analyses incorporating quadratic age terms demonstrated that the proposed framework remains applicable to more complex regression models accounting for non-linear aging trajectories. The spatial organization of significant effects and the relative behavior of the smoothing approaches remained globally consistent with the linear analyses. This suggests that the influence of smoothing strategies on statistical topology is robust across different model parameterizations and not restricted to linear aging effects alone.

Additionally, the effect of including the quadratic term in the nonlinear age model can be assessed by performing an F-test on the quadratic term to determine whether it captures additional variance. The results are presented in Figure 7, at the p<.001 uncorrected threshold for the sake of sensitivity. They reveal that the quadratic term captures some variance, at a uncorrected statistical threshold, in all quantitative maps. This demonstrates that incorporating the quadratic age term in the GLM can enhances the local interpretation of age-related effects. Therefore, accounting for a nonlinear age model would be recommended for precise quantification of age effects on the brain, particularly in studies aiming for fine-grained regional interpretations. However, for this methodological paper, we preferred to stick to the original, i.e. linear, model from Callaghan et al. for easier comparison and interpretability.

**Figure 7:**
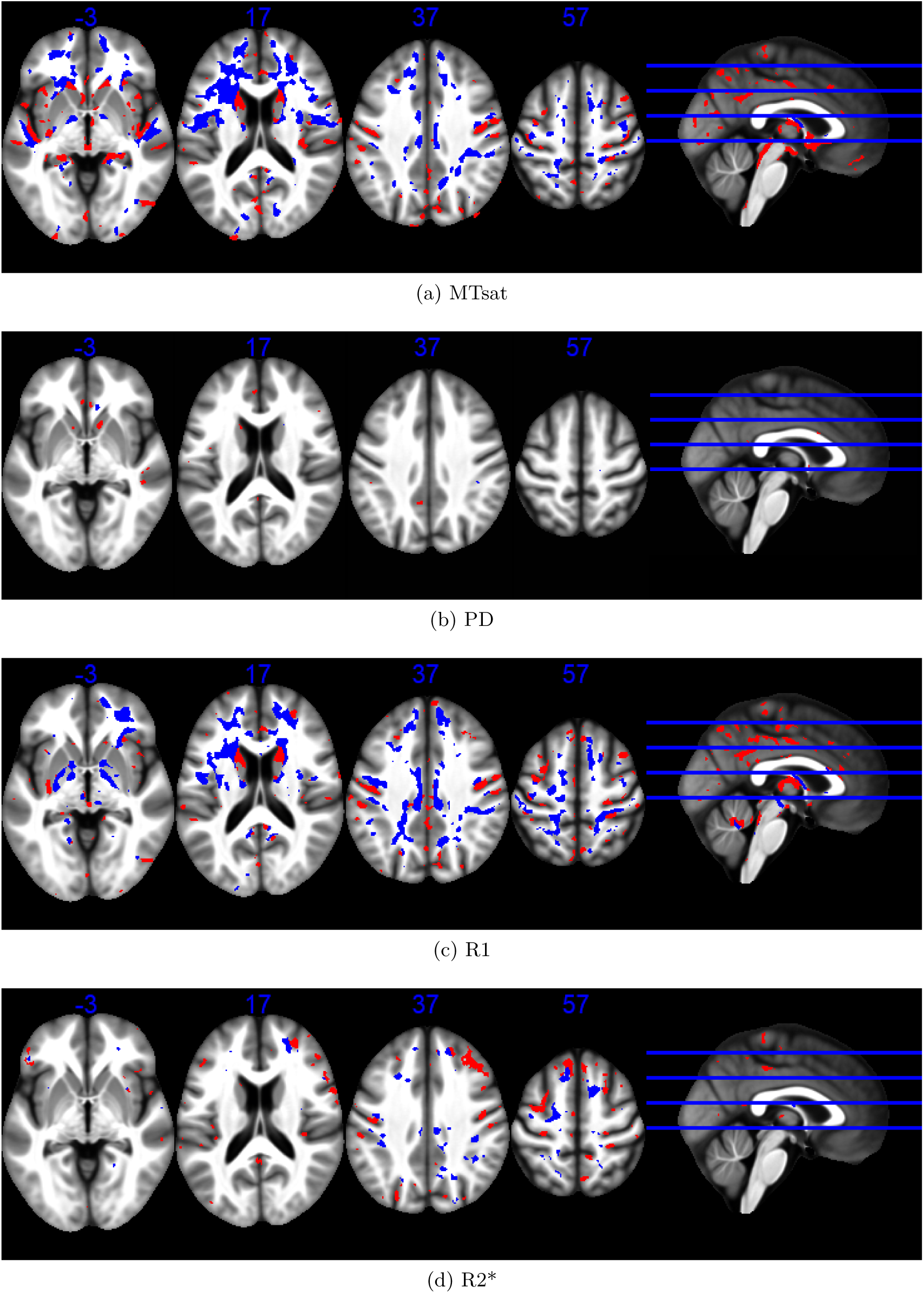
Statistical parametric maps identifying regions (red for GM and blue for WM) in which the quantitative parameters significantly varied with the quadratic age term at the *p <* 0.001 uncorrected level. The results are superimposed on the mean MT map for the cohort in MNI space. The four axial slices are located at z = −3, 17, 37 and 57 mm, from left to right, as illustrated on the sagital slice (right). SPM has been used for parametric statistical inference under stationarity assumption in RFT.

### **A.2** Effect of smoothing on thresholded SPMs

Figures 8, 9 and 10 illustrate representative voxel-wise statistical maps obtained for PD, R1 and R2* age-related effects using parametric inference under stationarity assumption. Figure 11 illustrates the TWS and gTSPOON MTsat statistical maps. The corresponding SUSANs map is in Figure 2.

**Figure 8:**
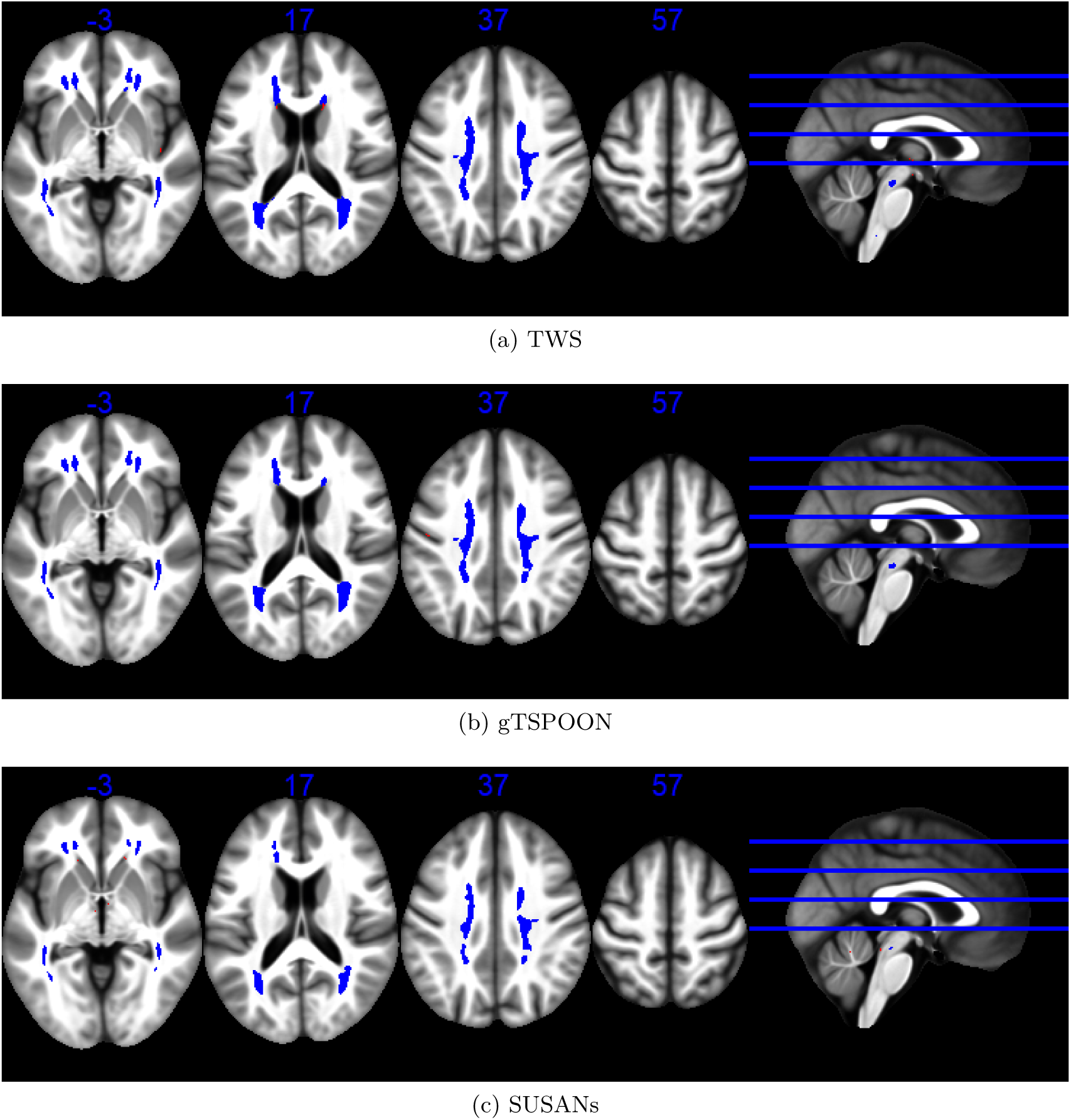
Statistical parametric maps identifying regions (red for GM and blue for WM) in which PD significantly increased with age at the *p <* 0.05 FWE corrected level. The results are superimposed on the mean MT map for the cohort in MNI space. The four axial slices are located at z =-3, 17, 37 and 57 mm, from left to right, as illustrated on the sagital slice (right). SPM has been used for parametric statistical inference under stationarity assumption in RFT.

**Figure 9:**
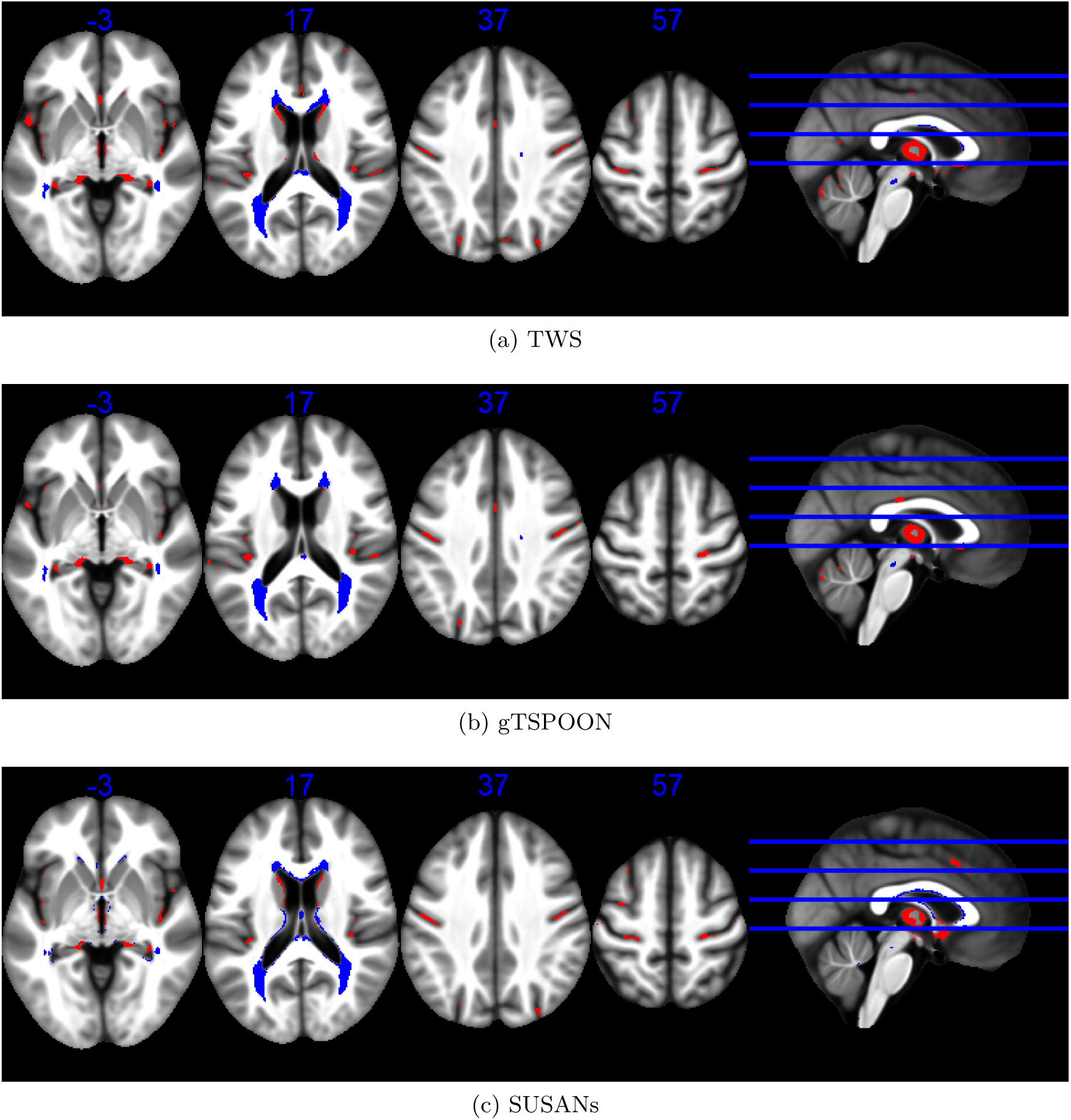
Statistical non-parametric maps identifying regions (red for GM and blue for WM) in which R1 significantly decreased with age at the *p <* 0.05 FWE corrected level. The results are superimposed on the mean MT map for the cohort in MNI space. The four axial slices are located at z =-3, 17, 37 and 57 mm, from left to right, as illustrated on the sagital slice (right). SPM has been used for parametric statistical inference under stationarity assumption in RFT.

**Figure 10:**
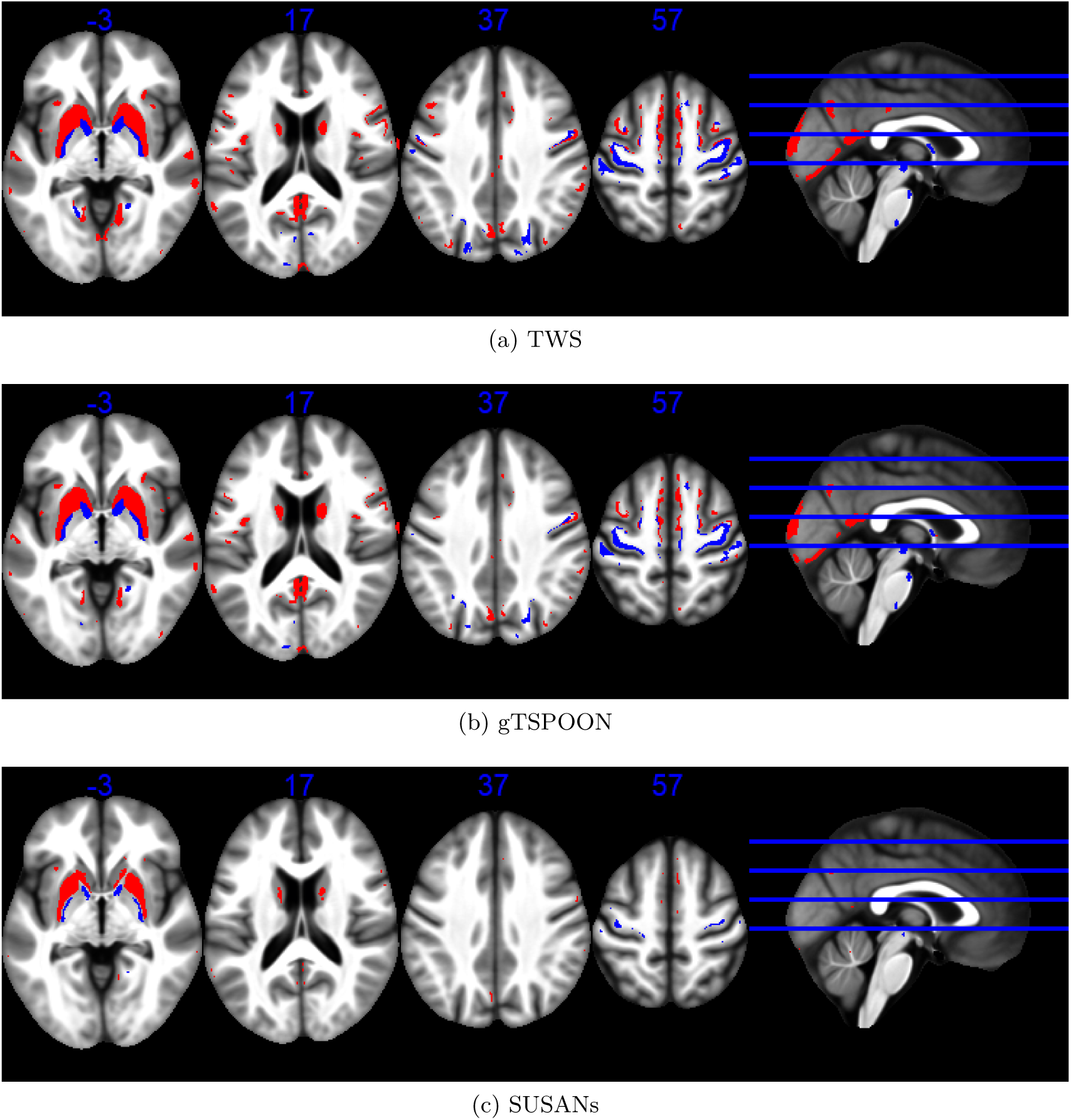
Statistical non-parametric maps identifying regions (red for GM and blue for WM) in which R2* significantly increased with age at the *p <* 0.05 FWE corrected level. The results are superimposed on the mean MT map for the cohort in MNI space. The four axial slices are located at z =-3, 17, 37 and 57 mm, from left to right, as illustrated on the sagital slice (right). SPM has been used for parametric statistical inference under stationarity assumption in RFT.

**Figure 11:**
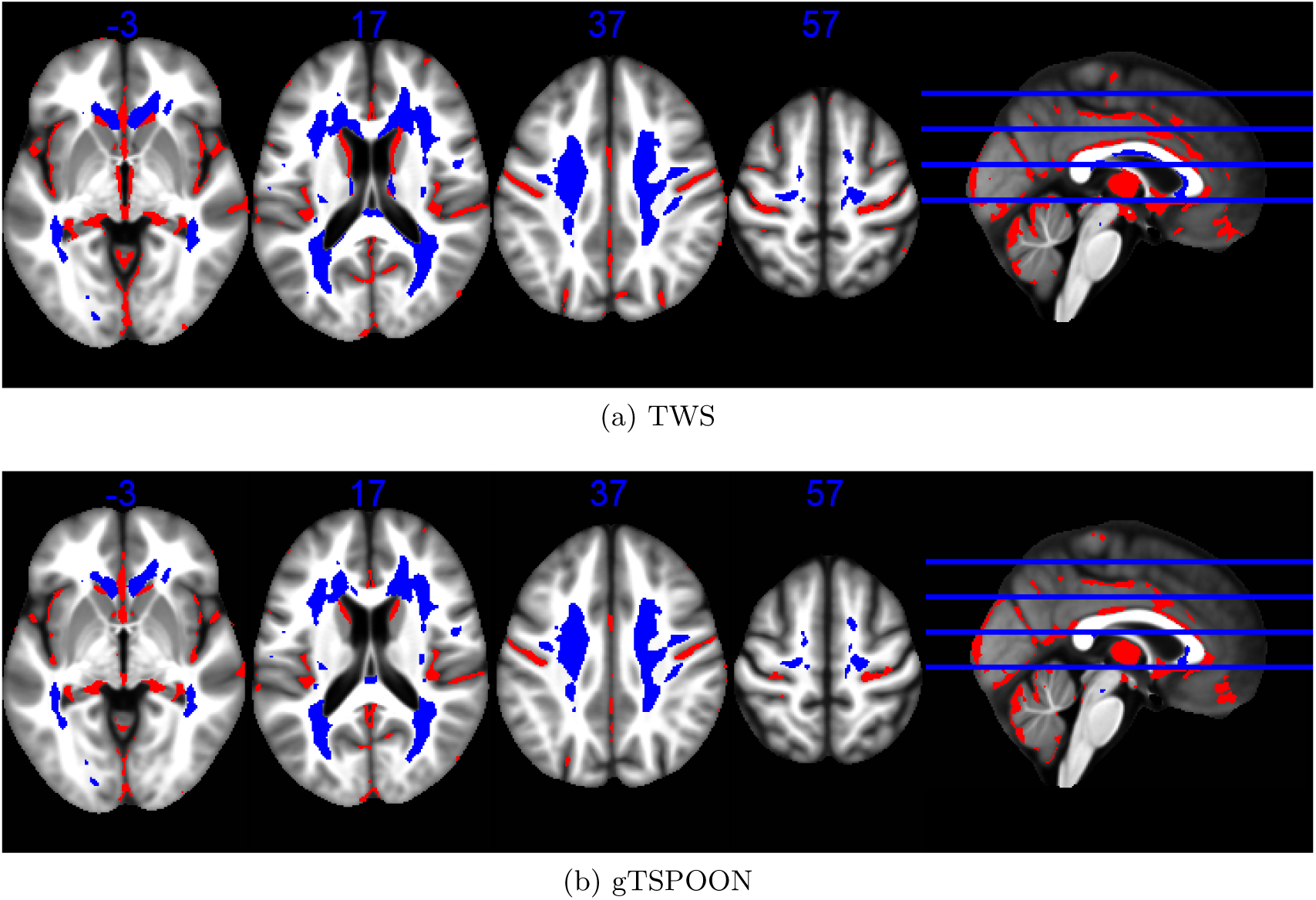
Statistical parametric maps identifying regions (red for GM and blue for WM) in which MTsat significantly decreased with age at the *p <* 0.05 FWE corrected level. The results are superimposed on the mean MT map for the cohort in MNI space. The four axial slices are located at z =-3, 17, 37 and 57 mm, from left to right, as illustrated on the sagital slice (right). SPM has been used for parametric statistical inference under stationarity assumption in RFT.

Figures 12, 13 and 14 illustrate representative voxel-wise statistical maps obtained for PD, R1 and R2* age-related effects using parametric inference under non stationarity assumption. Figure 15 illustrates the TWS and gTSPOON MTsat statistical maps. The corresponding SUSANs map is in Figure 2.

**Figure 12:**
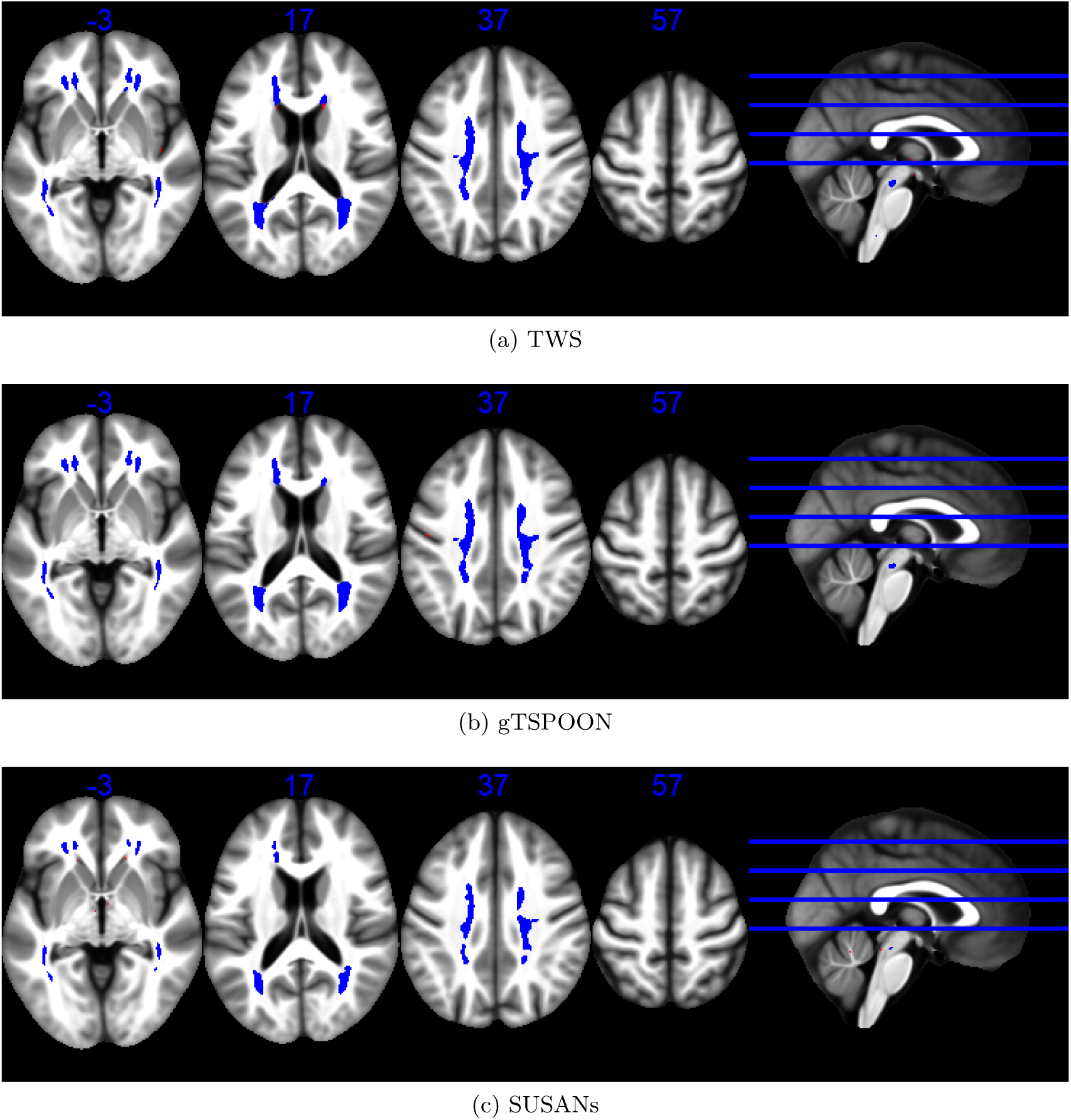
Statistical parametric maps identifying regions (red for GM and blue for WM) in which PD significantly increased with age at the *p <* 0.05 FWE corrected level. The results are superimposed on the mean MT map for the cohort in MNI space. The four axial slices are located at z =-3, 17, 37 and 57 mm, from left to right, as illustrated on the sagital slice (right). SPM has been used for parametric statistical inference under non stationarity assumption in RFT.

**Figure 13:**
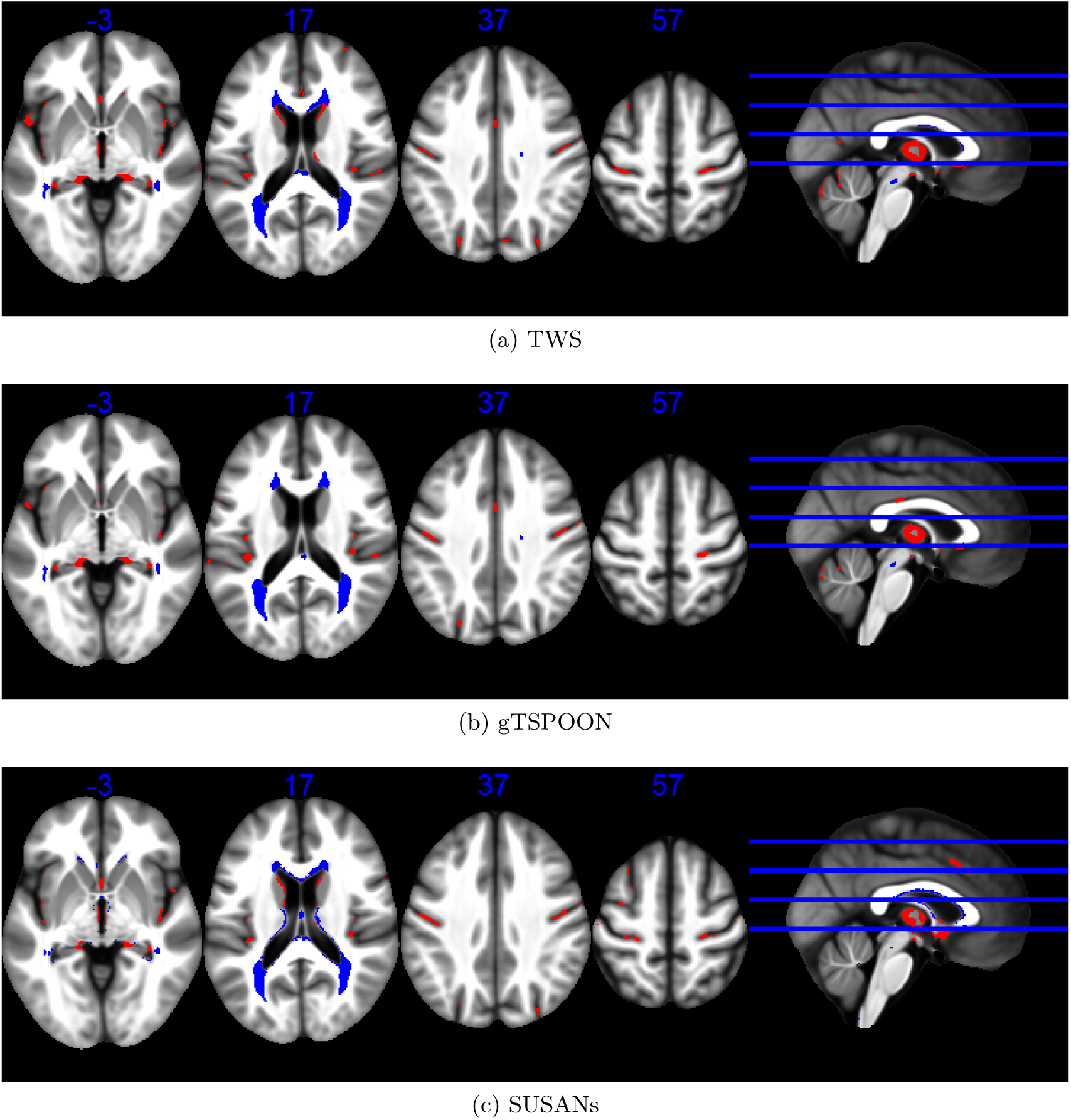
Statistical non-parametric maps identifying regions (red for GM and blue for WM) in which R1 significantly decreased with age at the *p <* 0.05 FWE corrected level. The results are superimposed on the mean MT map for the cohort in MNI space. The four axial slices are located at z = - 3, 17, 37 and 57 mm, from left to right, as illustrated on the sagital slice (right). SPM has been used for parametric statistical inference under non stationarity assumption in RFT.

**Figure 14:**
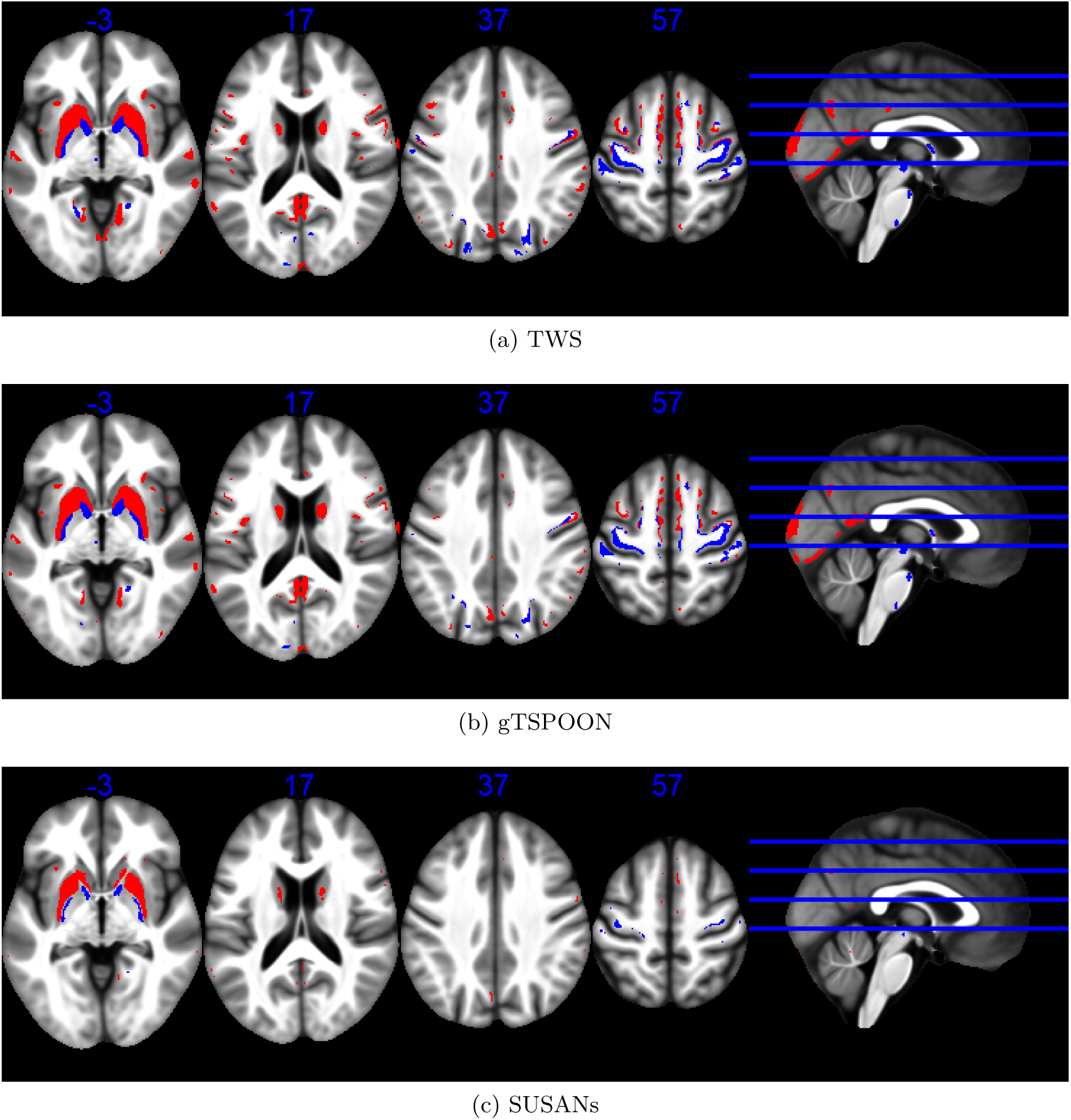
Statistical non-parametric maps identifying regions (red for GM and blue for WM) in which R2* significantly increased with age at the *p <* 0.05 FWE corrected level. The results are superimposed on the mean MT map for the cohort in MNI space. The four axial slices are located at z =-3, 17, 37 and 57 mm, from left to right, as illustrated on the sagital slice (right). SPM has been used for parametric statistical inference under non stationarity assumption in RFT.

**Figure 15:**
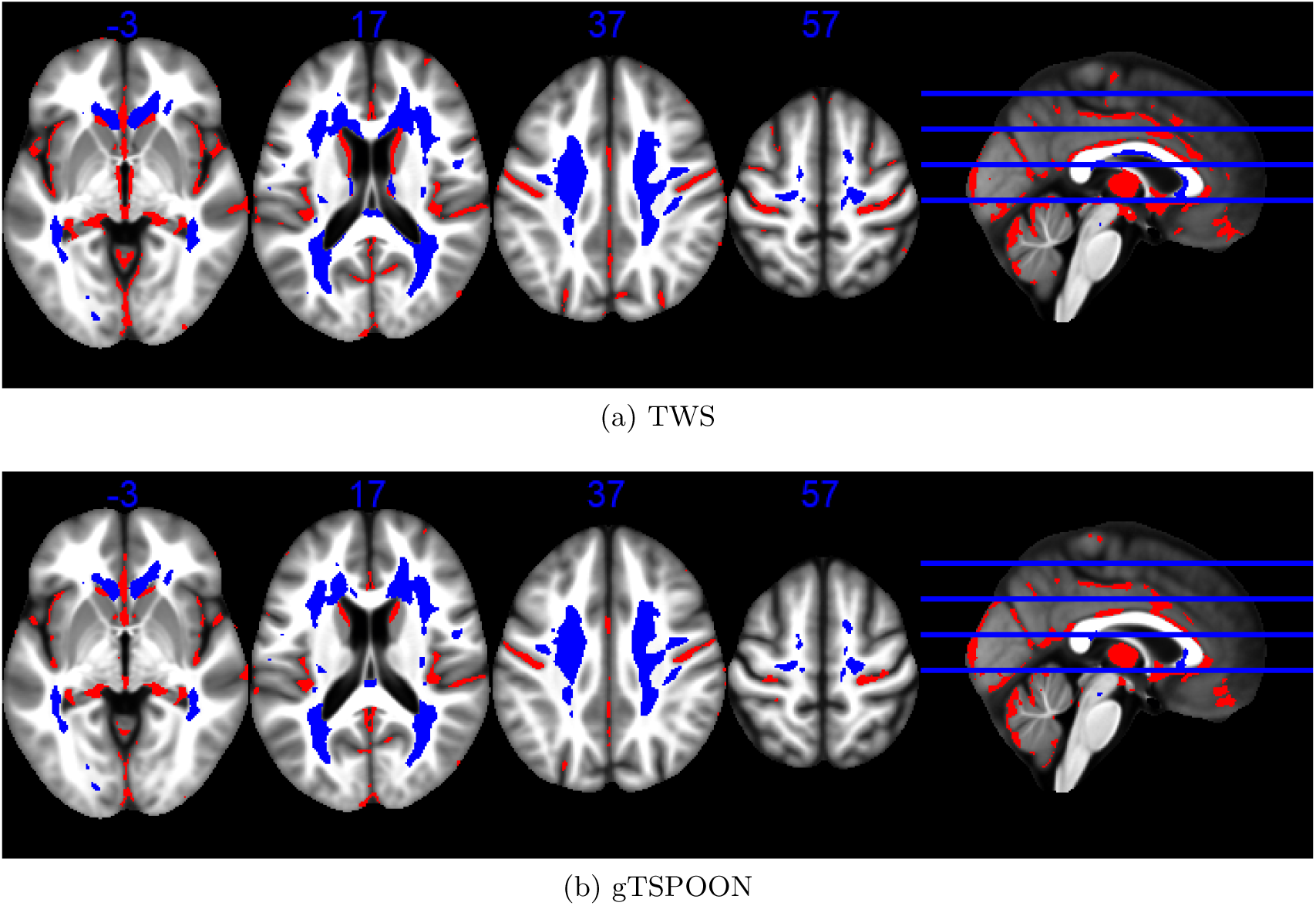
Statistical parametric maps identifying regions (red for GM and blue for WM) in which MTsat significantly decreased with age at the *p <* 0.05 FWE corrected level. The results are superimposed on the mean MT map for the cohort in MNI space. The four axial slices are located at z =-3, 17, 37 and 57 mm, from left to right, as illustrated on the sagital slice (right). SPM has been used for parametric statistical inference under non stationarity assumption in RFT.

Figures 16, 17 and 18 illustrates representative voxel-wise statistical maps obtained for PD, R1 and R2* age-related effects using permutation-based non parametric inference.

**Figure 16:**
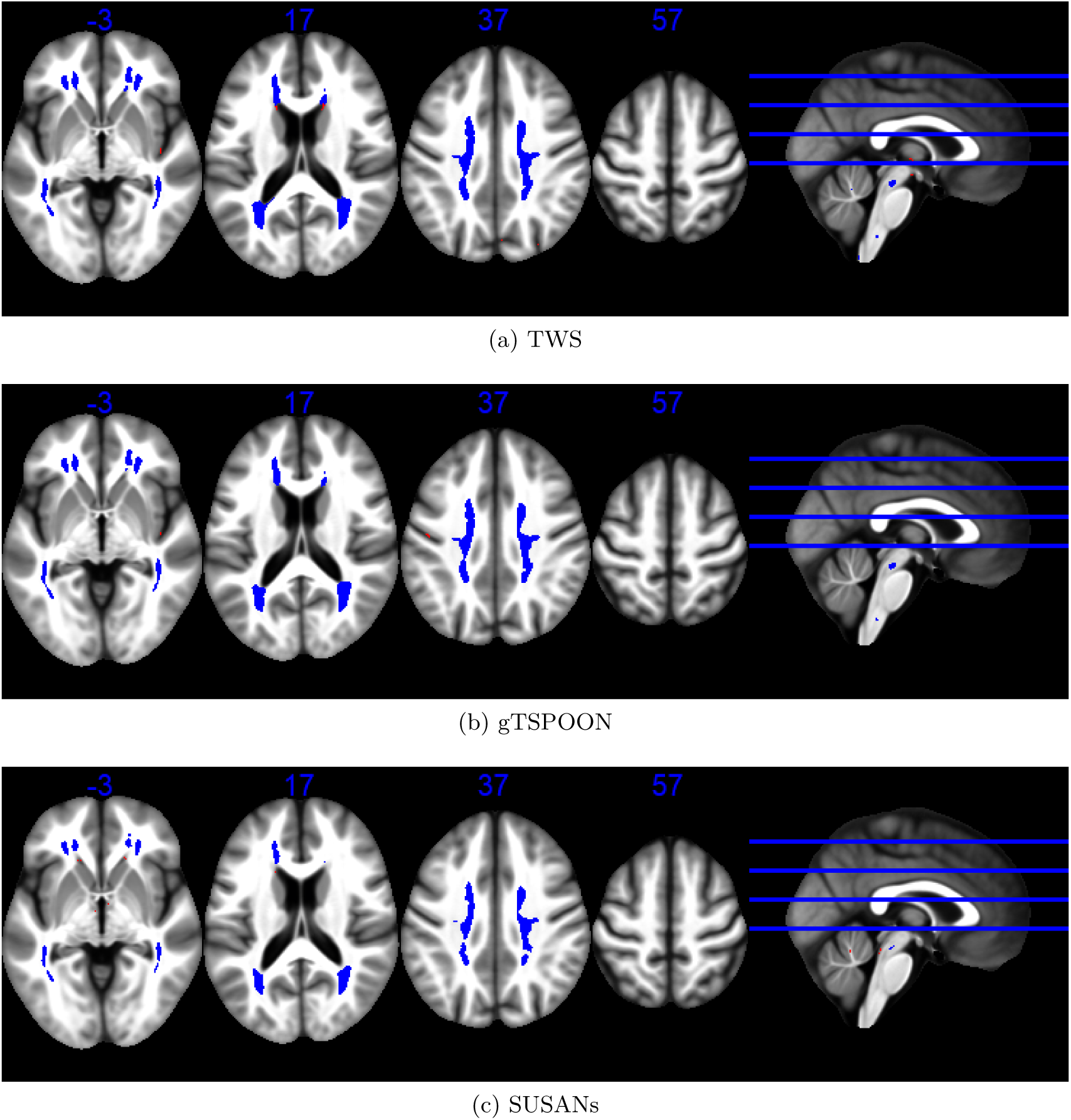
Statistical non-parametric maps identifying regions (red for GM and blue for WM) in which PD significantly increased with age at the *p <* 0.05 FWE corrected level. The results are superimposed on the mean MT map for the cohort in MNI space. The four axial slices are located at z =-3, 17, 37 and 57 mm, from left to right, as illustrated on the sagital slice (right). SnPM has been used for non parametric statistical inference (SnPM).

**Figure 17:**
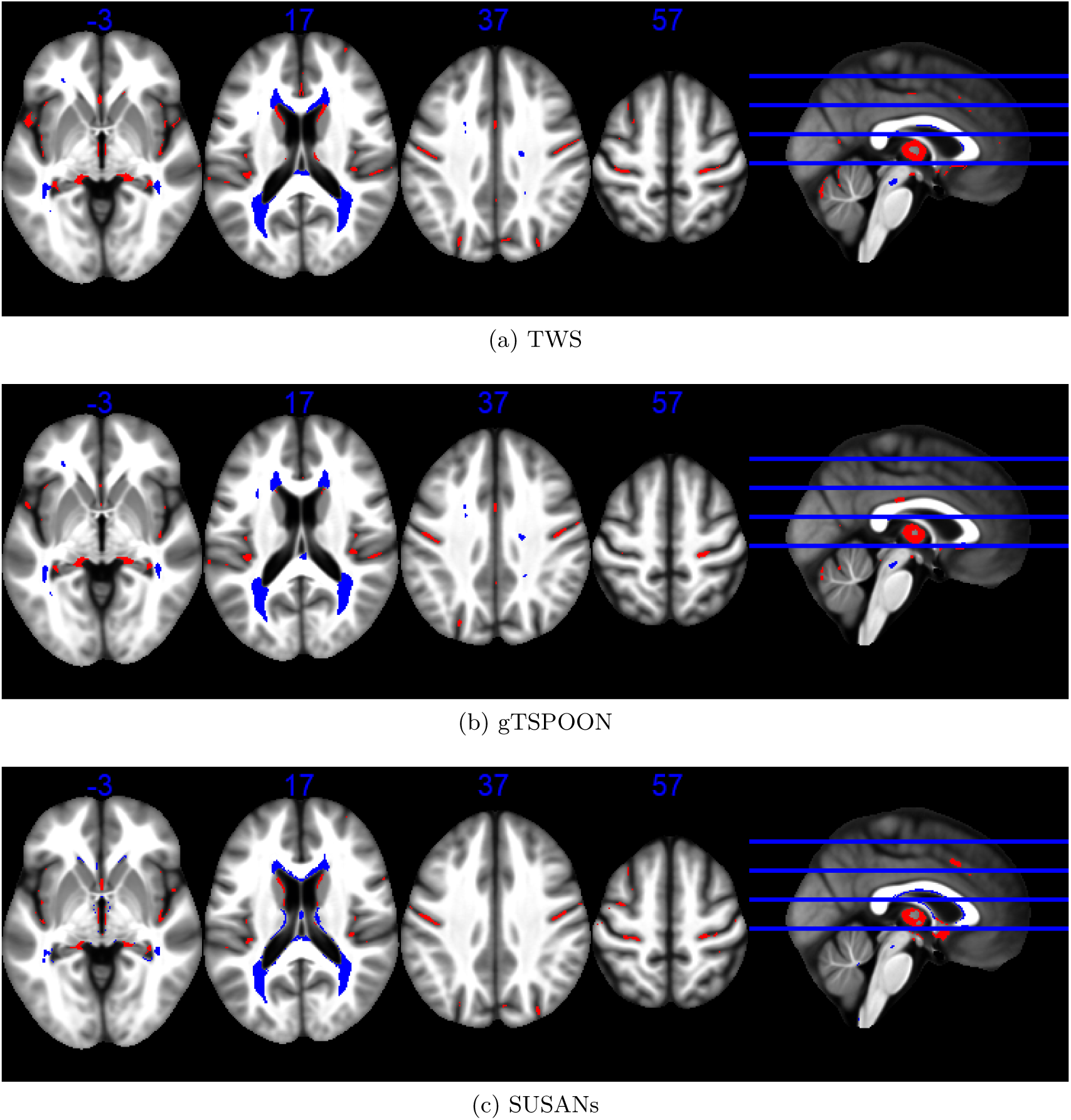
Statistical non-parametric maps identifying regions (red for GM and blue for WM) in which R1 significantly decreased with age at the *p <* 0.05 FWE corrected level. The results are superimposed on the mean MT map for the cohort in MNI space. The four axial slices are located at z =-3, 17, 37 and 57 mm, from left to right, as illustrated on the sagital slice (right). SnPM has been used for non parametric statistical inference (SnPM).

**Figure 18:**
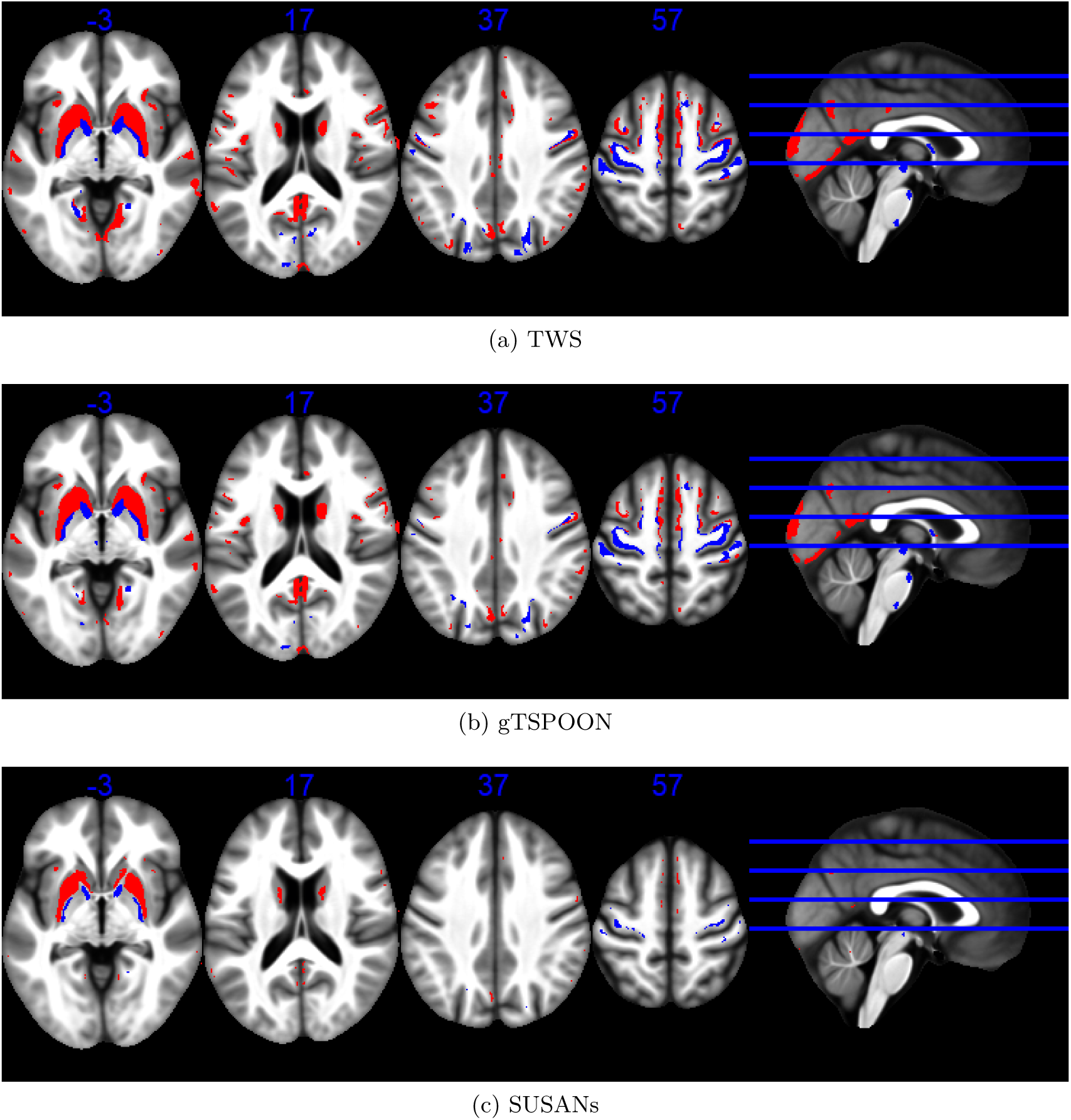
Statistical non-parametric maps identifying regions (red for GM and blue for WM) in which R2* significantly increased with age at the *p <* 0.05 FWE corrected level. The results are superimposed on the mean MT map for the cohort in MNI space. The four axial slices are located at z =-3, 17, 37 and 57 mm, from left to right, as illustrated on the sagital slice (right). SnPM has been used for non parametric statistical inference (SnPM).

The voxel-wise log-likelihood maps (Figs. 19, 20, 21) make it possible to highlight the minimization of residuals during GLM estimation as a function of the different smoothing approaches for PD, R1, and R2* parameters.

**Figure 19:**
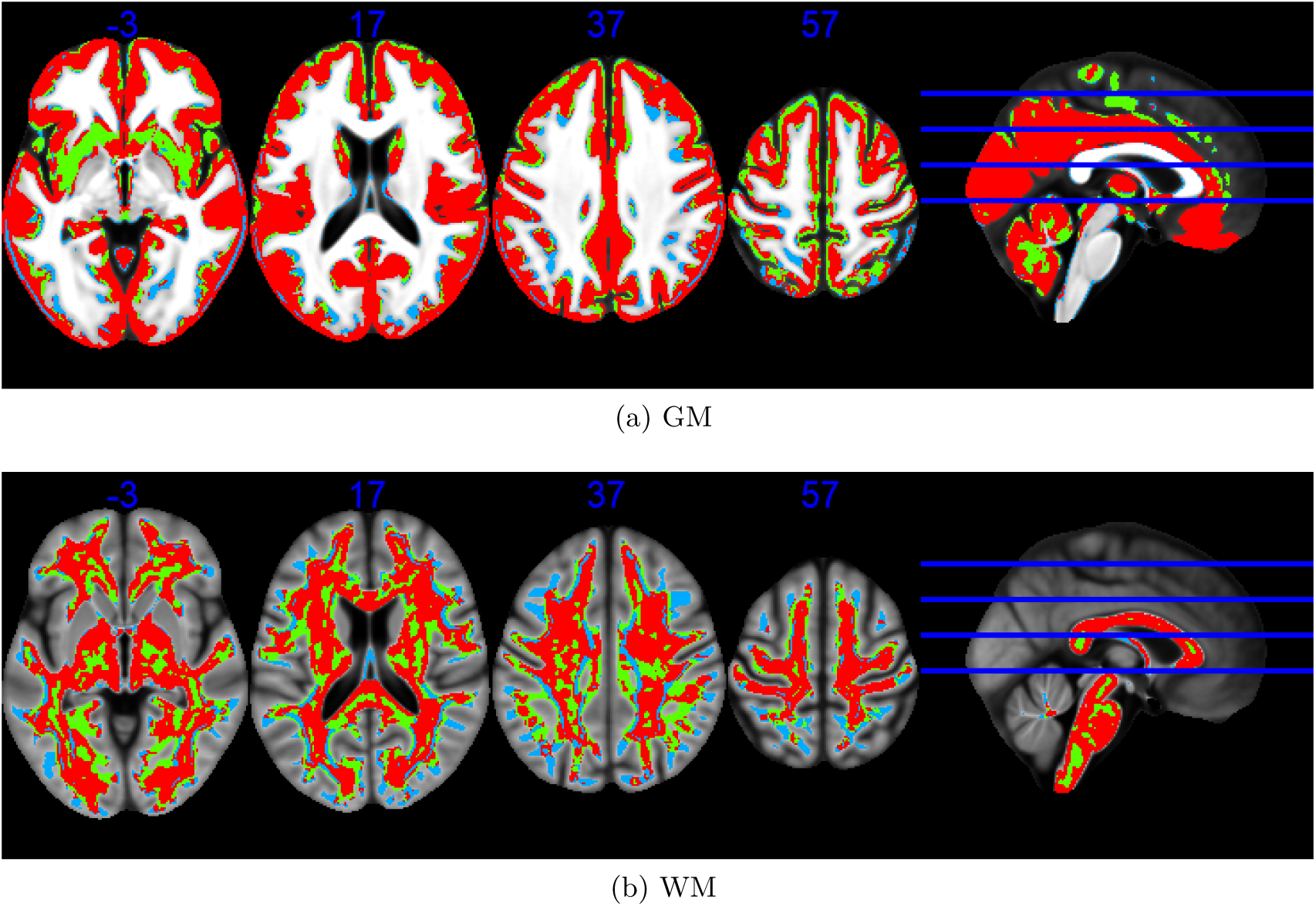
Voxelwise log-likelyhood maps (GM at the top and WM at the bottom) identifying regions in which GLM’s residuals are minimized on the PD maps smoothed using TWS, gTSPOON and SUSANs. Red regions shows best GLM fitting with the TWS-smoothed maps, green with gTSPOON-smoothed maps and blue with SUSANs-smoothed maps. The four axial slices are located at z = - 3, 17, 37 and 57 mm, from left to right, as illustrated on the sagital slice (right).

**Figure 20:**
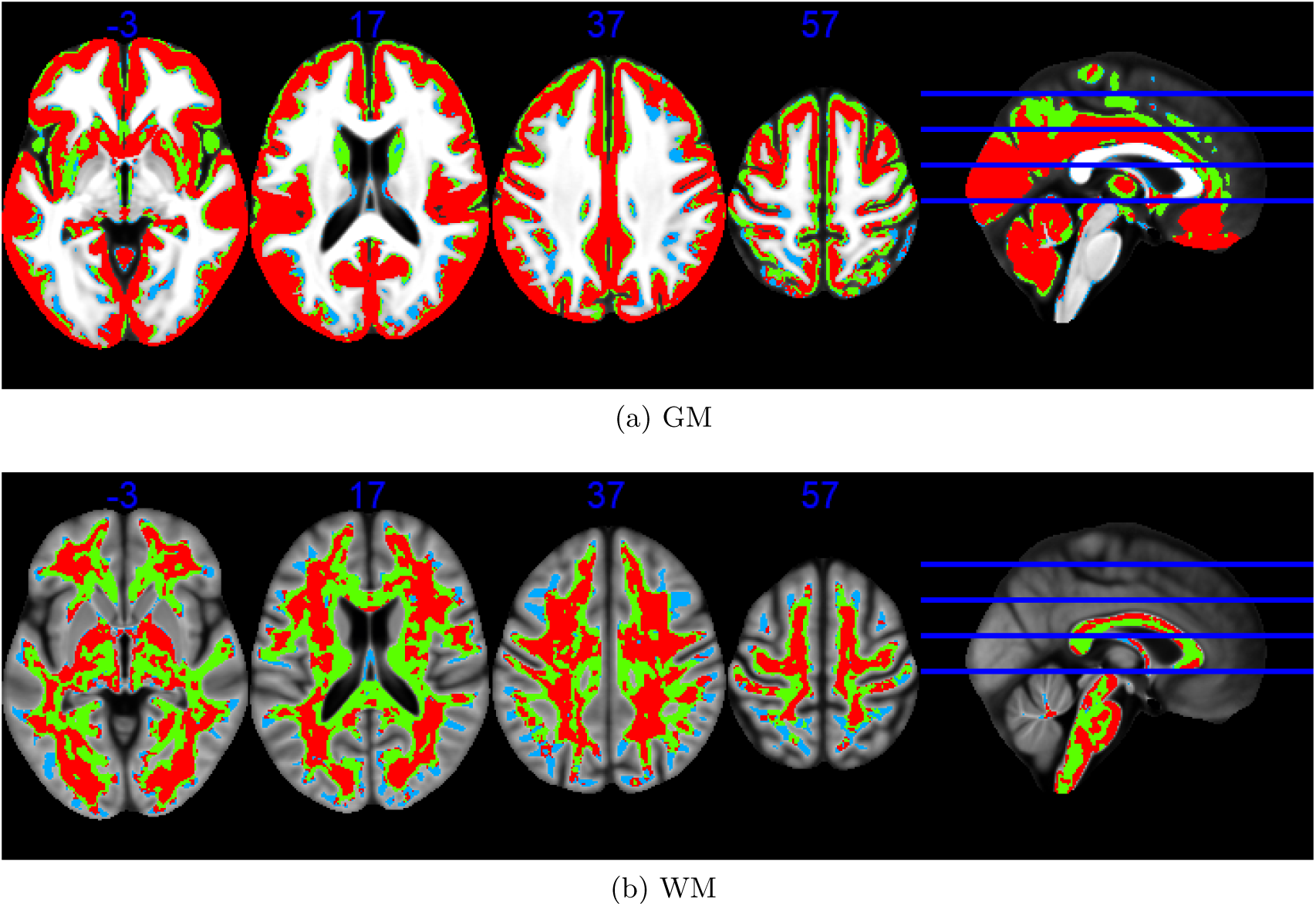
Voxelwise log-likelyhood maps (GM at the top and WM at the bottom) identifying regions in which GLM’s residuals are minimized on the R1 maps smoothed using TWS, gTSPOON and SUSANs. Red regions shows best GLM fitting with the TWS-smoothed maps, green with gTSPOON-smoothed maps and blue with SUSANs-smoothed maps. The four axial slices are located at z = - 3, 17, 37 and 57 mm, from left to right, as illustrated on the sagital slice (right).

**Figure 21:**
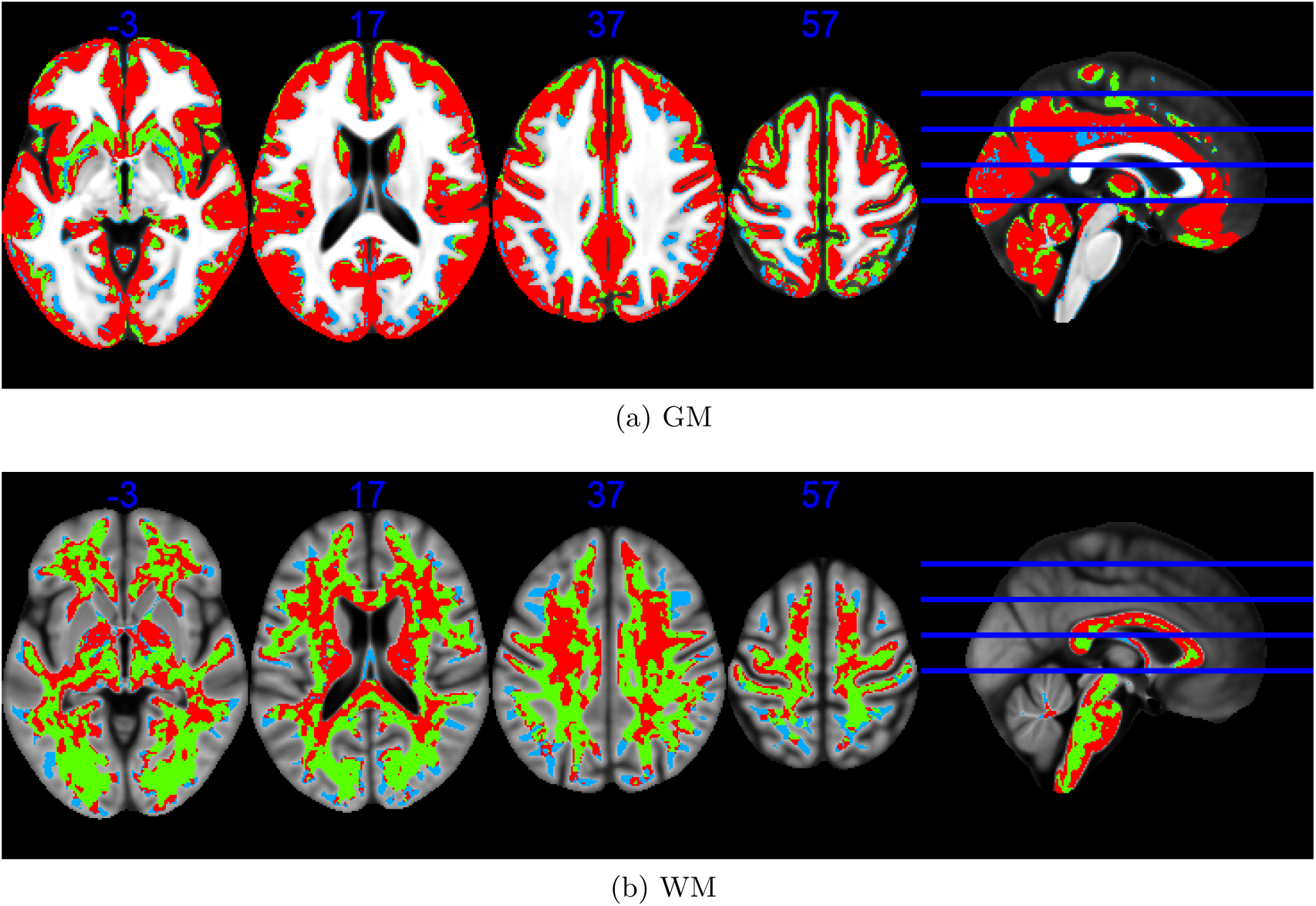
Voxelwise log-likelyhood maps (GM at the top and WM at the bottom) identifying regions in which GLM’s residuals are minimized on the R2* maps smoothed using TWS, gTSPOON and SUSANs. Red regions shows best GLM fitting with the TWS-smoothed maps, green with gTSPOON-smoothed maps and blue with SUSANs-smoothed maps. The four axial slices are located at z = - 3, 17, 37 and 57 mm, from left to right, as illustrated on the sagital slice (right).

## Notes

https://doi.org/10.18112/openneuro.ds005851.v1.0.0

### Competing Interest Statement

The authors have declared no competing interest.

